# Developmental isoform diversity in the human neocortex informs neuropsychiatric risk mechanisms

**DOI:** 10.1101/2023.03.25.534016

**Authors:** Ashok Patowary, Pan Zhang, Connor Jops, Celine K. Vuong, Xinzhu Ge, Kangcheng Hou, Minsoo Kim, Naihua Gong, Michael Margolis, Daniel Vo, Xusheng Wang, Chunyu Liu, Bogdan Pasaniuc, Jingyi Jessica Li, Michael J. Gandal, Luis de la Torre-Ubieta

## Abstract

RNA splicing is highly prevalent in the brain and has strong links to neuropsychiatric disorders, yet the role of cell-type-specific splicing or transcript-isoform diversity during human brain development has not been systematically investigated. Here, we leveraged single-molecule long-read sequencing to deeply profile the full-length transcriptome of the germinal zone (GZ) and cortical plate (CP) regions of the developing human neocortex at tissue and single-cell resolution. We identified 214,516 unique isoforms, of which 72.6% are novel (unannotated in Gencode-v33), and uncovered a substantial contribution of transcript-isoform diversity, regulated by RNA binding proteins, in defining cellular identity in the developing neocortex. We leveraged this comprehensive isoform-centric gene annotation to re-prioritize thousands of rare de novo risk variants and elucidate genetic risk mechanisms for neuropsychiatric disorders.

**One-Sentence Summary:** A cell-specific atlas of gene isoform expression helps shape our understanding of brain development and disease.

**Structured Abstract:** *INTRODUCTION:* The development of the human brain is regulated by precise molecular and genetic mechanisms driving spatio-temporal and cell-type-specific transcript expression programs. Alternative splicing, a major mechanism increasing transcript diversity, is highly prevalent in the human brain, influences many aspects of brain development, and has strong links to neuropsychiatric disorders. Despite this, the cell-type-specific transcript-isoform diversity of the developing human brain has not been systematically investigated.

*RATIONALE:* Understanding splicing patterns and isoform diversity across the developing neocortex has translational relevance and can elucidate genetic risk mechanisms in neurodevelopmental disorders. However, short-read sequencing, the prevalent technology for transcriptome profiling, is not well suited to capturing alternative splicing and isoform diversity. To address this, we employed third-generation long-read sequencing, which enables capture and sequencing of complete individual RNA molecules, to deeply profile the full-length transcriptome of the germinal zone (GZ) and cortical plate (CP) regions of the developing human neocortex at tissue and single-cell resolution.

*RESULTS:* We profiled microdissected GZ and CP regions of post-conception week (PCW) 15-17 human neocortex in bulk and at single-cell resolution across six subjects using high-fidelity long-read sequencing (PacBio IsoSeq). We identified 214,516 unique isoforms, of which 72.6% were novel (unannotated in Gencode), and >7,000 novel exons, expanding the proteome by 92,422 putative proteoforms. We uncovered thousands of isoform switches during cortical neurogenesis predicted to impact RNA regulatory domains or protein structure and implicating previously uncharacterized RNA-binding proteins in cellular identity and neuropsychiatric disease. At the single-cell level, early-stage excitatory neurons exhibited the greatest isoform diversity, and isoform-centric single-cell clustering led to the identification of previously uncharacterized cell states. We systematically assessed the contribution of transcriptomic features, and localized cell and spatio-temporal transcript expression signatures across neuropsychiatric disorders, revealing predominant enrichments in dynamic isoform expression and utilization patterns and that the number and complexity of isoforms per gene is strongly predictive of disease. Leveraging this resource, we re-prioritized thousands of rare de novo risk variants associated with autism spectrum disorders (ASD), intellectual disability (ID), and neurodevelopmental disorders (NDDs), more broadly, to potentially more severe consequences and revealed a larger proportion of cryptic splice variants with the expanded transcriptome annotation provided in this study.

*CONCLUSION:* Our study offers a comprehensive landscape of isoform diversity in the human neocortex during development. This extensive cataloging of novel isoforms and splicing events sheds light on the underlying mechanisms of neurodevelopmental disorders and presents an opportunity to explore rare genetic variants linked to these conditions. The implications of our findings extend beyond fundamental neuroscience, as they provide crucial insights into the molecular basis of developmental brain disorders and pave the way for targeted therapeutic interventions. To facilitate exploration of this dataset we developed an online portal (https://sciso.gandallab.org/).

## Main Text

Human brain development is a tightly coordinated process under precise molecular and genetic control. Transcriptomics has provided substantial insights into the cellular and molecular processes regulating human brain development (*1*, *2*), including characterization of the underlying heterogeneous cell-types, states, and lineages through single-cell RNA-sequencing (scRNA-Seq) (*3–5*) as well as their developmental trajectories (*6*), gene regulatory networks, and cell-to-cell variability (*7*). However, the technological limitations of short-read scRNA-Seq have largely prevented systematic characterization of the full complexity of cell-type-specific alternative splicing (AS) and resulting isoform diversity present during neurodevelopment (*8*).

AS is a fundamental form of tissue-specific gene regulation present in >90% of multi-exon genes (*9*). RNA transcript diversity results from the combinatorial effects of alternative transcription start sites (TSS), regulated exon inclusion/exclusion or intron retention, and distinct transcript termination sites due to alternative polyadenylation (APA) (*10*). In individual cases, synaptic genes such as *Nrxn1* have thousands of unique isoforms (*11*). Human brain-expressed genes, which are longer and contain the most exons, undergo the greatest degree of splicing compared with other tissues and species - a mechanism contributing to the vast proteomic, phenotypic, and evolutionary complexity of the human brain (*12*, *13*). Indeed, AS plays an important role in synaptogenesis, synapse specification (*14*) and brain development more broadly (*15*). Substantial cell-type-specificity of splicing and isoform diversity has been observed in the mouse brain, even among closely related neurons, and often with precise temporal regulation (*16–20*).

AS has been implicated as a critical mechanism linking genetic variation and neuropsychiatric disease (*2*, *21–24*). For example, recent work identified marked splicing and isoform expression dysregulation in the brains of individuals with ASD and schizophrenia (SCZ) - a signal substantially more widespread than gene expression changes with greater enrichment for genetic risk (*25–27*). Common variants associated with ASD and SCZ from genome-wide association studies (GWAS) showed substantial enrichment among splicing-QTLs in the developing and adult human brain (*2*, *24*). Likewise, rare de novo variants identified in ASD and ID are enriched for predicted splice-altering consequences (*23*). These genetic signals also exhibited convergence during mid-fetal brain development, highlighting the clinical relevance of this critical developmental period characterized by rapid increases in neurogenesis (*28*, *29*). Finally, a comprehensive understanding of splicing and isoform complexity can have direct therapeutic relevance, as demonstrated recently for spinal muscular atrophy (*30*).

The advent of third-generation, long-read sequencing technologies has enabled accurate, in-depth characterization of the full-length alternatively-spliced transcriptome at scale (*10*, *31*). Coupled with single-cell barcoding, recent work has begun to catalog the isoform-centric transcriptome with single-cell resolution (*20*, *32*). Here, we leverage this approach to deeply profile the major cell-types of the developing human neocortex at mid-gestation. We uncover >150k previously unannotated transcript-isoforms and thousands of spliced exons, many within high-confidence NDD risk genes. To functionally annotate these isoforms, we integrated proteomics, characterized regional patterns of isoform-switching during corticogenesis, embedded isoforms within coexpression networks, and mapped transcripts to 16 distinct cell-type clusters. Altogether, results highlight the tremendous complexity of transcript-isoform diversity during neurodevelopment, which we leverage to uncover mechanisms of cell-fate specification and genetic risk mechanisms for NDDs.

## Results

### Full-length transcriptome of the developing human brain

We performed high-depth, high-fidelity long-read sequencing (PacBio HiFi IsoSeq) to comprehensively profile the full-length polyadenylated transcriptome of the developing human neocortex across six donors at mid-gestation (15-17 PCW) (**Fig. 1A**). To interrogate patterns of differential transcript expression and usage during peak neurogenesis, samples were first microdissected into the neural progenitor-enriched germinal zone (GZ) and the neuron-enriched cortical plate (CP). Following sequencing and comprehensive quality control (**Fig. 1B-C**), we generated >33 million high-quality (>Q20) circular consensus sequence (CCS) reads across samples (**fig. S1-2**). Using minimap2 (*33*), >99% of full-length reads were confidently aligned to the reference genome, a marked improvement over the ∼85% mapping rate of short-read RNA-seq (**fig. S2A-B**). Using the TALON pipeline (*34*), 214,516 unique transcripts were found in the bulk tissue transcriptome (**data S1**), corresponding to 24,554 genes; of these, >175k isoforms from 17,299 genes were expressed at >0.1 transcripts per million (TPM) in ≥50% of the samples (**Fig. 1D-E**). Isoform-level expression exhibited reproducibility across technical and biological replicates (**Fig. 1B** and **fig. S2C**), with clear separation of CP and GZ samples (**Fig. 1C**). The median read length was 2.99 kb (range: 80-14200 bp), consistent with the high RNA quality and expected distribution of mammalian mRNA transcripts (**Fig. 1F**).

**Fig. 1.**
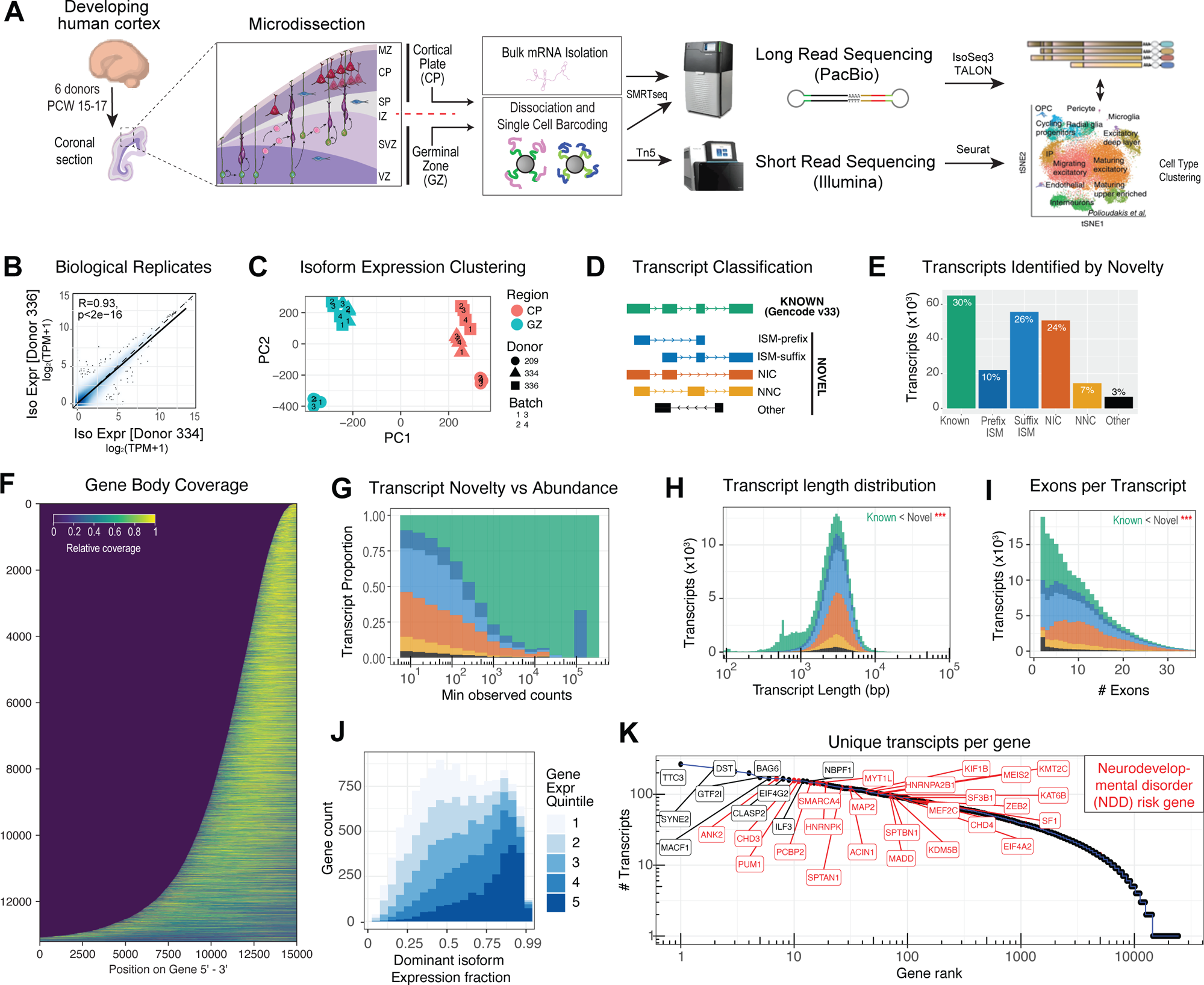
The full-length cell-type specific transcriptome of the developing human neocortex at mid-gestation. (**A**) Experimental design for isoform-centric profiling of the developing human brain transcriptome at bulk and single-cell resolution. Briefly, microdissected samples from the progenitor-enriched germinal zone (GZ) and neuronally-enriched cortical plate (CP) were profiled from 6 unique donors at mid-gestation. Full-length cDNA libraries were generated from homogenate tissue as well as from dissociated, barcoded single-cells with incorporation of UMIs. Single molecule long-read sequencing (PacBio) was used to quantify transcript-isoforms and integrated with matched short-read scRNA-Seq. Isoform expression quantifications demonstrated (**B**) strong biological reproducibility across unique donors and (**C**) region-specific clustering via principal component analysis. (**D**) Transcript isoforms identified by IsoSeq were compared against the Gencode v33 reference. Novel transcripts were further classified by their splice junction matching to annotated Gencode isoforms as described by TALON: ISM (incomplete splice match); NIC (novel in catalog); NNC (novel not in catalog). The ‘other’ category denotes isoforms belonging to antisense, genomic, and intergenic classes. (**E**) Number of isoforms identified based on classes described in D. (**F**) Heatmap shows uniform patterns of relative read-depth coverage across genes, arranged by length. Low coverage in dark blue, high coverage in yellow. (**G**) Abundance of the isoforms by each class as described in D. The Prefix ISM signal observed at Min observed counts = ∼10^5^ largely corresponds to a highly expressed isoform of the MAP1B gene. Compared with known isoforms, novel transcripts identified here were (**H**) significantly longer (P<2e^−16^, Kruskal-Wallis) and (**I**) contained a significantly greater number of exons (P<2e^−16^, Kruskal-Wallis). (**J**) Proportion of the dominant isoforms for each gene by gene expression percentile. For highly expressed genes, the dominant isoform contributed the most to the gene expression. (**K**) Genes ranked by the number of unique transcript isoforms detected. NDD risk genes (red) (*82*) had significantly more detected isoforms, controlling for total expression, gene length, and coding length (OR 1.56, P=3.6e-03, logistic regression).

We next generated a high-quality, custom reference annotation of the developing neocortex transcriptome (**table S1**), merging data across samples. Compared with Gencode v33, only 65,006 (30.3%) of observed isoforms matched existing transcripts (**Fig. 1D-E**). We further classified isoforms based on their splice junction match to Gencode, strand specificity, and 5’ and/or 3’ overlap with known transcripts (*35*) (**fig. 1D**; **table S1**). As a class, novel transcripts were more lowly expressed than known transcripts (**Fig. 1G**), although several were individually highly expressed, including *WASF1*, *CYFIP2*, *MAP1B*, *NEFL*, and *SMARCA4* (**fig. S3A-E**).

Most isoforms not found in Gencode were classified as incomplete splice match (ISM; **fig 1D-E**). Although ISM transcripts are often disregarded as artifacts of RNA degradation or internal priming, gene body coverage did not show evidence of 3’ bias (**Fig. 1F; fig. S2A**). Further, novel transcripts (including ISMs) were significantly longer (P<2e-16, Kruskal-Wallis; **Fig. 1H**) and contained more exons (P<2e-16, Kruskal-Wallis; **Fig. 1I**) compared with known transcripts. Consequently, we retained ISM transcripts with additional supporting evidence (*36*), and note that many exhibited functional roles as “hub” isoforms in network analyses detailed below.

Finally, given this large number of isoforms detected, we sought to characterize their patterns of usage and potential functional relevance to NDDs. For multi-isoform genes, we calculated the proportion of gene expression attributable to the top (dominant) isoform, stratified by total gene abundance (**Fig. 1J**). Whereas lowly expressed genes exhibited a relatively even distribution, for highly expressed genes, the dominant isoform tended to capture the majority of expressed reads. Nevertheless, the number of detected isoforms per gene was strongly predictive of NDD risk gene status, even accounting for gene length and total gene expression (P=5.2e-03, logistic regression; **Fig. 1K**). This association remained significant even when restricting only to novel transcripts (OR 1.8, P=5.5e-04, **fig. S3F**).

### Expanded transcriptomic and proteomic complexity in the developing human brain

To contextualize this expanded transcriptomic complexity in the developing brain, we next integrated several orthogonal genomic annotations (*37–43*) and predicted potential downstream protein coding consequences (**Fig. 2**). Altogether, ∼80% of novel isoform TSS’s were supported by proximal, annotated CAGE and/or ATAC-Seq peaks. At the 3’ end, ∼91% of novel transcripts were supported by nearby polyA sites/motifs–a rate higher than for known transcripts (**Fig. 2A-B**). Of 38,115 splice junctions not observed in Gencode, 74% were validated by Intropolis (**Fig. 2C**). Finally, 53% of novel transcripts were validated when combining the latest version of Gencode (v43) with multiple independent long-read datasets, despite the majority representing non-neural tissues (**fig. S3G**). Together, these results provided broad orthogonal support for novel isoforms discovered in this study.

**Fig. 2.**
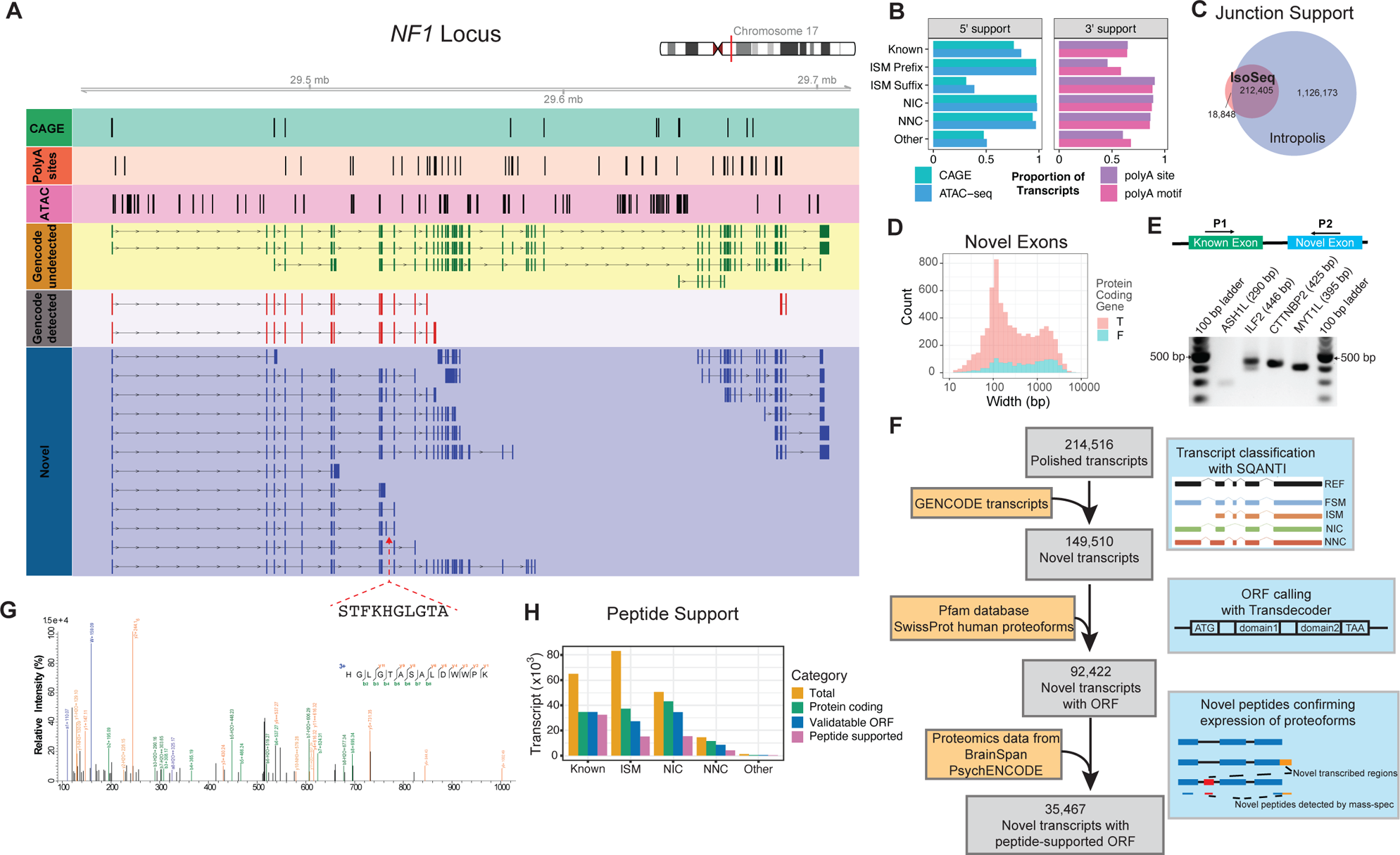
Expanded Transcriptomic and Proteomic Complexity. (**A**) The *NF1* gene locus with 20 previously unidentified brain-expressed isoforms. Tracks from top to bottom include: CAGE clusters, 3′seq clusters, ATAC-Seq peaks, Gencode isoforms that were not detected in our data, Gencode isoforms detected in our data, and novel isoforms identified from our data. A novel microexon is highlighted. (**B**) External validation of isoforms by independent datasets: 5’ end validation was performed by presence of peak from CAGE (FANTOM5 and fetal brain cortex) and mid-gestation cortex ATAC-Seq (*37–39*); 3’ end validation was performed by presence of polyA motif or peak from polyAsite database. Percentage of transcript with support at either end is highlighted. (**C**) The vast majority of splice junctions identified by IsoSeq are supported by the external Intropolis splice junction database. (**D**) Length distribution of the >7000 novel spliced exons uncovered in this study. T=True, F= False. (**E**) Validation of novel exons via RT-PCR. Expected product size is shown in parentheses along the name of the gene with the exon. Each exon was amplified with the primer sets shown in the schematic. (**F**) Characterization of novel protein-coding transcripts. Long-read sequencing identified a total of 214,516 transcripts, 149,510 of which were not found in Gencode v33. 92,422 of these novel transcripts were predicted to code for protein sequences, and 35,467 predicted ORFs were further confirmed by MS/MS proteomics data. (**G**) Representative mass spectrum of peptide HGLGTASALDWWPK, which confirms the translation of the identified *NF1* microexon. Matched b, y, a, and immonium ions were highlighted. (**H**) Number of total transcripts, transcripts with ORF, transcripts with novel ORF compared to UniProt human protein sequences, and transcripts with ORFs validated by MS/MS proteomics, plotted per isoform structural category.

In total, compared with Gencode, our data extended by ∼27Mb the transcribed portion of the human genome. Of this, 3.85 Mb comprised >7000 previously unannotated spliced exons, spanning >3500 unique genes (**Fig. 2D**). Nearly 80% of these exons were supported by canonical splice junctions (**table S2**) and we successfully validated several using RT-PCR (**Fig. 2E**; **table S2**). Additionally, we identified 319 multi-exonic genes not matching any existing gene model (**table S1**), of which 256 were antisense to existing genes and 63 were located within intergenic regions. For example, we identified an 18-exon gene with 4 unique splice isoforms antisense to the ASD risk gene *DPYSL2* (**fig. S3H**).

We next sought to determine the functional relevance of this expanded compendium of novel exons and transcripts at the protein-coding level (**Fig. 2F**). Of the novel transcripts observed, 92,422 exhibited protein coding potential, with at least one complete putative open reading frame (ORF). Integration of human brain proteomics data (*44*) provided peptide-level support for 35,467 novel transcripts with unique proteoforms (**table S2**). For example, a novel, alternatively spliced 30 bp microexon found in the NDD risk gene *NF1* was predicted to add an additional 10 amino acids (AA) to the known protein (**Fig. 2A**). Searching against mass-spectrometry proteomics data, we identified spectra (n=15) that confidently supported the existence of this 10 AA sequence (**Fig. 2G**). Notably, isoforms containing this microexon were only detected in neuronally-enriched CP samples (**fig. S4A**). Extending these analyses beyond a single gene locus, we observed broad peptide-level support for novel transcripts across all classification categories (**Fig. 2H**).

### Isoform Switching during Neurogenesis

We next sought to contrast gene and isoform usage across GZ/CP samples to identify genes exhibiting isoform-switching during neurogenesis and/or neuronal maturation. Whereas differential gene expression (DGE) signatures of neuronal maturation have been extensively characterized in the developing human brain at bulk tissue and single-cell resolution (*3*, *4*, *37*), AS and isoform usage have not been investigated transcriptome-wide during this critical developmental period.

Consistent with previous work (*3*), many genes exhibited significant DGE patterns between GZ and CP (4,475 of 24,554 genes, at FDR <0.05; **table S3**). Likewise, of genes with multiple expressed isoforms, a large proportion exhibited differential transcript usage (DTU) across GZ and CP (2,679 of 10,809 genes at FDR <0.05; **Fig. 3A**; **table S3**), with the majority (57%) of significantly switching isoforms (5,630 isoforms at FDR<0.05) coming from novel transcripts (**Fig. 3B**). Although there was a significant overlap among DGE and DTU genes (1,010 genes, OR=2, P<10^−46^, Fisher’s exact test; **Fig. 3A**), 1,669 genes exhibited isoform switching without changes in overall expression. The majority of GZ/CP isoform switching events had observed and/or predicted functional consequences (**Fig. 3C**), summarized in **table S3**. Of 5,630 isoform switching events, 3,204 (57%) were predicted to alter the ORF length, with longer ORFs predicted among CP-upregulated isoforms (FDR-corrected P<0.0001). However, CP-upregulated isoforms were also more likely to exhibit predicted nonsense-mediated decay (NMD) sensitivity (243 vs 100 isoforms; FDR-corrected P<10^−15^).

**Fig. 3.**
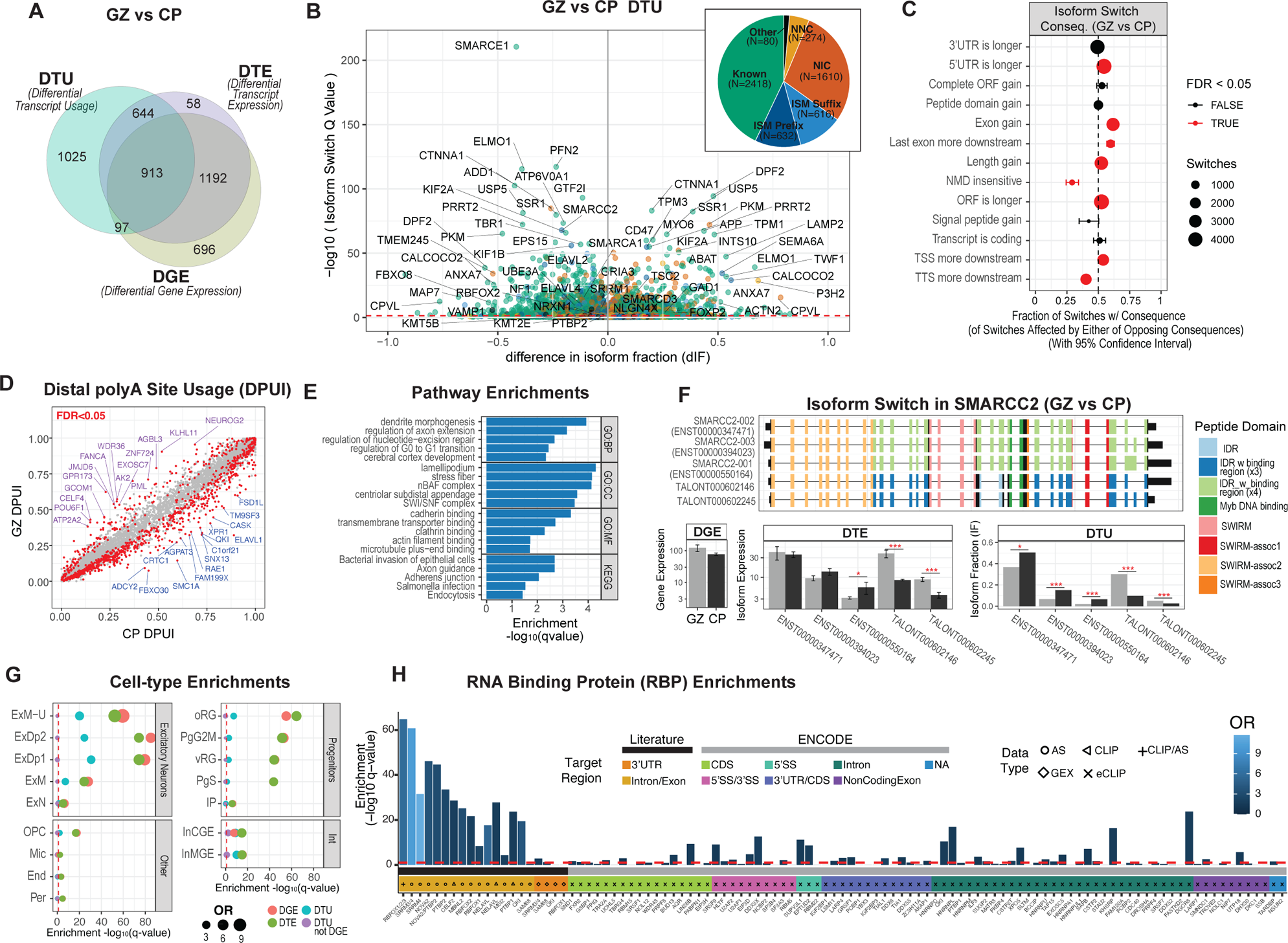
The landscape of isoform switching during human corticogenesis. (**A**) Long-read RNA-Seq data from GZ and CP samples were contrasted for patterns of differential gene expression (DGE), differential transcript expression (DTE), and differential transcript usage (DTU). Venn diagram is shown depicting the overlap for genes exhibiting significant DGE, DTE, and DTU (FDR-corrected P<0.05). (**B**) A volcano plot depicts isoform switching across GZ and CP. The x-axis depicts the difference in isoform fraction (dIF) for a given transcript in CP versus GZ. Inset, most regionally variable DTU isoforms are not present in Gencode. (**C**) Functional consequences of isoform switch events between GZ and CP are shown. For example, CP-upregulated isoforms were more significantly likely to have gained (than lost) an exon (2,031 vs 1,260 isoforms, FDR-corrected P<10^−40^). (**D**) Analysis of distal polyA site usage index (DPUI) for known transcripts between GZ and CP samples. On average, CP transcripts have higher DPUIs indicative of longer 3’ UTRs. (**E**) Pathway enrichments for genes exhibiting cross-region DTU are notable for dendrite morphogenesis and SWI/SNF complex genes, among others. (**F**) An example of isoform switching observed within the ASD risk gene *SMARCC2*. Although total gene expression was not different between GZ and CP, significant switching was observed among DTU isoforms with two isoforms exhibiting preferential usage in GZ (* P<0.05, *** P<0.001). (**G**) Regionally variable genes were enriched for cell-type specific marker genes from scRNA-Seq. vRG = ventricular radial glia, oRG = Outer Radial Glia, PgG2M = Cycling progenitors (G2/M phase), PgS = Cycling progenitors (S phase), IP = Intermediate Progenitors, ExN = Migrating excitatory, ExM = Maturing excitatory, ExM-U = Maturing excitatory upper enriched, ExDp1 = Excitatory deep layer 1, ExDp2 = Excitatory deep layer 2, InMGE = Interneuron MGE, InCGE = Interneuron CGE, OPC = Oligodendrocytes Precursor Cells, End = Endothelial, Per = Pericyte, Mic = Microglia. (**H**) Genes containing DTU isoforms were also highly enriched for targets of known brain-enriched RBPs (black bar) and for targets of RBPs profiled in the ENCODE database (gray bar). Targets exhibiting alternative splicing (AS) or gene expression (GEX), or direct binding (e/CLIP) are indicated by Data Type. See also **fig. S4C**.

To further interrogate how the 3’ untranslated region (UTR) of transcripts differed between regions, we performed a complementary, read-depth based analysis of distal polyadenylation (polyA) usage (**Fig. 3D**). For the 9,896 transcripts with multiple annotated polyA sites, we used DaPars2 to compute a distal polyA usage index (DPUI) - the fraction of total reads mapping to the longer (distal) 3’UTR (*45*). In total, 1,013 transcripts exhibited significant differences in DPUI between GZ and CP (repeated measures ANOVA; FDR-corrected P < 0.05; **Fig. 3D; table S3**). DPUIs were increased in CP versus GZ for the majority (772) of transcripts, and DPUI was on average greater in the CP (two-sample paired T-test, p < 2.2e-16), indicating overall increased 3’UTR lengths with neuronal maturation. Pathway enrichment among genes exhibiting GZ/CP DPUI differences was notable for RNA/mRNA binding as well as cytoplasmic stress granule gene ontologies. Finally, we found RNA-binding proteins (RBPs) (*46*) were overrepresented among transcripts exhibiting significant DPUI changes (453/3527 RBP genes vs 560/6369 non-RBP genes; one-tailed Fisher’s Exact Test, p = 2.06e-10). These findings indicated that 3’UTR lengthening in the CP is prevalent among genes encoding RBPs, suggesting that tight regulation of RBP activity is important for the transition from neural progenitor to neuron.

We next conducted pathway analyses for all genes exhibiting significant (DTU, one-sided Fisher’s exact test, P<0.05, **table S3A**) isoform switches across regions. Enriched biological pathways included dendrite morphogenesis, cadherin binding, and the chromatin modifier nBAF and SWI/SNF complexes, known to harbor convergent genetic risk for neuropsychiatric disorders (**Fig. 3E**; (*47*, *48*)). For example, among the top genes was *SMARCC2*, a high-confidence ASD risk gene encoding a subunit of the SWI/SNF chromatin remodeling complex (**Fig. 3F**). Although overall expression of *SMARCC2* did not differ between CP and GZ, two newly identified isoforms of *SMARCC2* showed preferential usage in GZ and exhibited exon skipping compared with known transcripts. Other notable isoform-switching genes included several known splicing regulators and RBPs (*SRRM1*, *SRRM4, CELF1, PTBP2*, *ELAVL2*, *ELAVL4*, *RBFOX2*), chromatin modifiers (*KMT2E*, *KMT5B*, *SMARCA1, SMARCD3*, *SMARCE1*), transcription factors (*FOXP2*), regulators of synaptic transmission (*GRIA3*, *VAMP1*, *GAD1*), and synaptic adhesion molecules (*NLGN4X*, *NRXN1*). Isoform-switching genes were broadly expressed across cell-types, with particular enrichment for excitatory neuron lineages (**Fig. 3G**).

### Putative RNA-binding protein regulators of isoform switching events

RBPs are a diverse class of proteins regulating the processing and fate of target mRNAs. Through regulation of AS, RBPs alter isoform and consequently protein diversity and play important roles in mammalian neural development and function (*17*, *49*). To identify potential RBP regulators of isoform switching events in the developing human brain, we assessed DGE, DTE, and DTU genes for enrichment for RBP targets identified through AS changes or direct binding by cross-linking immunoprecipitation (CLIP) using two datasets – 1) targets of RBPs that regulate AS during neural development and/or maturation curated from previous work (“brain-enriched”; black bar, **Fig. 3H and fig. S4B-C**), and 2) targets defined by systematic profiling of RBPs in the ENCODE repository (gray bar; **Fig. 3H and fig. S4B-C**).

Examining isoform diversity, we found that DTU genes significantly overlapped with targets of known regulators of AS in the brain (one-sided Fisher’s exact test, FDR-corrected P < 0.05, **Fig. 3H**; (*49*)). Splicing targets of brain-enriched RBPs are more enriched in DTU genes than DTE/DGE genes – consistent with known roles of these RBPs as regulators of AS in the brain (*49*). Correspondingly, gene expression targets, rather than AS targets, were more enriched in DGE and DTE than DTU genes (**fig. S4B**). We also observed that gene expression targets of RBPs that are more highly expressed in neurons than progenitors (*SRRM3/4*; (*49*)) are enriched in genes that increase in expression over neural development, while those that are targets of progenitor-associated RBPs (*SAM68*, *PTBP1*; (*49*)) show the opposite trend (**fig. S4C**).

To identify RBPs important for but not previously studied during the developmental transition from progenitor to neuron, we performed enrichment analysis of DTU, DTE, and DGE genes compared to a comprehensive set of RBP targets compiled from ENCODE (**Fig. 3H**; (*50*)). Similar to our results comparing target genes of brain-enriched RBPs, we found that ENCODE RBP targets were more enriched among DTU genes compared to DTE/DGE. Within DTU genes, ENCODE RBPs targeting introns, 5’/3’ splice sites, and 3’ splice sites were more enriched than those targeting the 3’UTR/CDS or 3’UTRs (**Fig. 3H**; dark green, magenta, and aqua boxes vs. green and dark blue boxes). This result matches the overall trend for RBP target enrichment in DTU over DGE, the latter of which is often mechanistically regulated through 3’ UTR binding.

Of the ENCODE RBPs showing significant target overlap with DTU genes (one-sided Fisher’s exact test, FDR-corrected P < 0.05, **Fig. 3H**), several have been increasingly recognized to play critical roles in neural development and disease. These included RBPs known to regulate RNA metabolism and splicing such as *LIN28B*, *EFTUD2*, *KHSRP*, and *DGCR8* (*51–53*). We observed a strong enrichment between DTU genes and targets of *DDX3X*, an X-linked RNA helicase where de novo mutations lead to sexually dimorphic ID and ASD (*54*, *55*). We also found strong enrichment of DTU genes among RBP targets with known roles in RNA metabolism, but which have not yet been studied in the context of neural development. For example, targets bound by components of the exosome (the RNA degradation system) and those involved in rRNA biogenesis, such as *EXOSC5*, *UTP18*, and *SUPV3L1* (*56–58*), and the nuclear matrix protein *SAFB*, implicated in heterochromatin regulation (*59*), are enriched in DTU genes. Altogether, these results indicate that while many of the GZ/CP isoform switching events are likely regulated by brain-enriched RBPs through AS, many more switching events are expected to be produced through diverse mechanisms regulated by RBPs previously not known to function in neural development.

### Network Context of Developmental Isoform Regulation

Given the large number of isoform switching events, we next leveraged weighted gene correlation network analysis (WGCNA) to place these results within a systems-level context during human brain development (*60*, *61*). We separately built unsupervised co-expression networks for gene (geneExpr), isoform expression (isoExpr), and transcript usage quantifications (isoUsage), the proportion of each gene’s total abundance attributable to a given isoform. For each network, genes or isoforms were assigned to modules based on shared patterns of covariation across samples (**fig. S5**; **table S4**), enabling in silico deconvolution of cell-type isoform usage as well as functional inference through pathway analysis (**fig. S5**).

We next compared properties across the three networks to better understand which factors drove differential transcript/gene expression and transcript usage. We observed that while both geneExpr and isoExpr networks were strongly driven by cell-type identity, isoExpr modules were enriched for specific progenitor and neuron subtypes (**Fig. 4A** and **fig. S6**). Multiple modules across isoExpr/isoUsage networks were enriched for RNA processing, cytoskeletal function and chromatin regulation pathways (**fig. S7A-D**). We further identified several disease-associated modules including isoExpr.M11, where a novel isoform of the ASD risk gene *DDX3X* is a hub transcript (**fig. S7E**). Together, these results show that isoform-level transcript expression further refines the resolution of cell type-specific modules and that isoform regulation is important for neurogenesis.

**Fig. 4.**
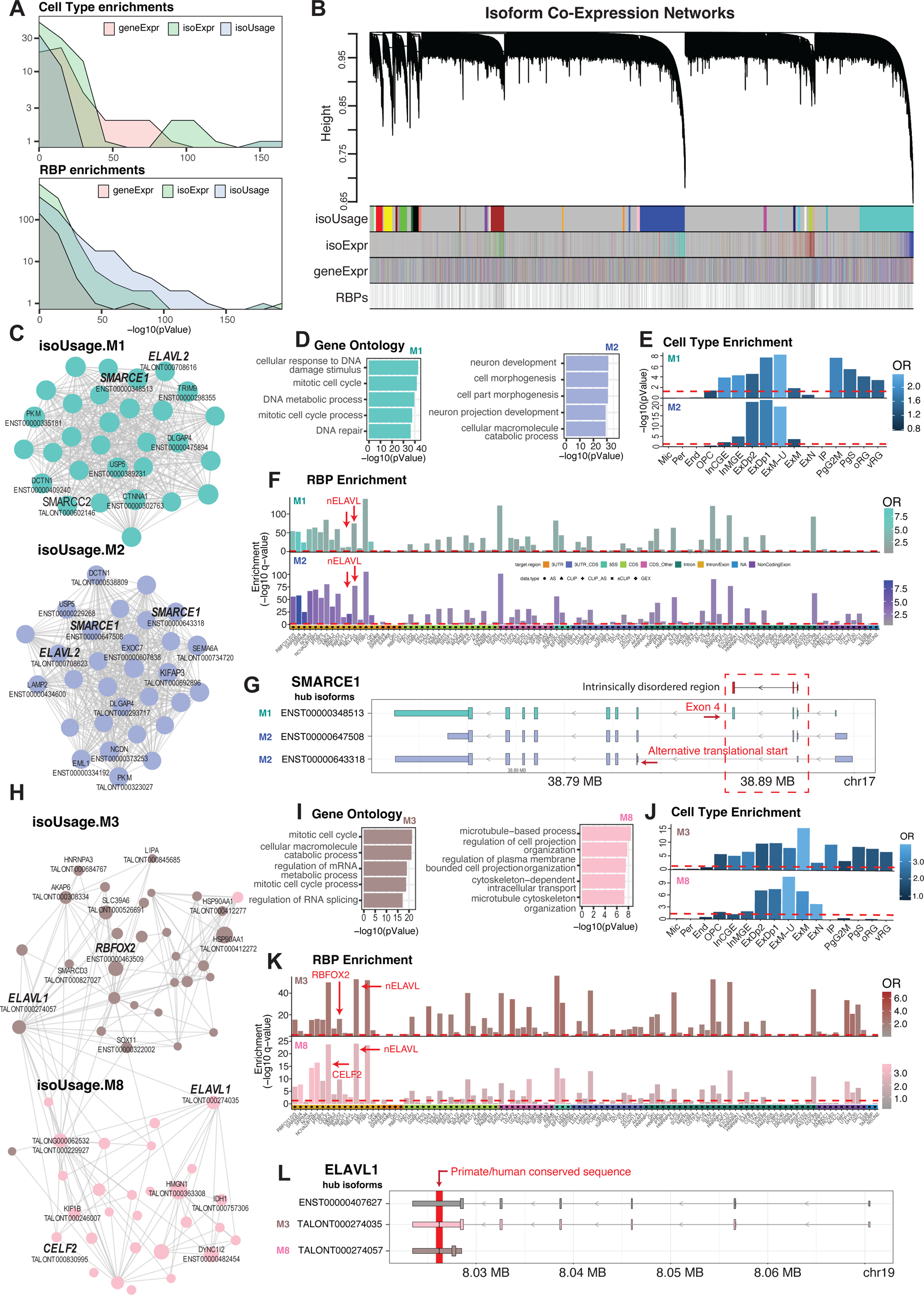
Network-based contextualization of isoform usage. (**A**) The isoUsage network shows more and stronger enrichments for RBP targets compared to geneExpr and isoExpr. Top, density plot of cell type enrichments for the three networks; bottom, density plot of RBP enrichments. (**B**) The isoUsage network is driven by RNA-binding protein isoform usage. Dendrogram of the isoUsage network with isoforms (isoUsage and isoExpr) or genes (geneExpr) organized by their presence in isoUsage modules plotted below. Isoforms of known RBPs (*112*) are plotted below. (**C**) Module plots highlighting hub isoforms in isoUsage.M1 and M2. *SMARCE1* hub isoforms inform different cellular processes associated with progenitors and neurons. (**D**) Gene ontology for iso.Usage.M1 and M2. (**E**) Cell type marker enrichment for iso.Usage.M1 and M2. (**F**) RBP target enrichments for M1 and M2, including targets of the nELAVL RBPs, which include *ELAVL1*. (**G**) Transcript models of *SMARCE1* hub isoforms. Box highlights exon3 (M1, turquoise) which encodes part of the IDR and the shifted reading frame driven by an alternative translational start (M2, blue). M2 SMARCE1 isoforms lack either all or a portion of this IDR – in ENST00000647508, exclusion of exon 4 truncates the IDR, while in ENST00000643318 a downstream translational start in combination with exon 4 exclusion entirely removes the protein domain. (**H**) Module plots for iso.Usage.M3 and M8. (**I**) Gene ontology for M3 and M8. (**J**) Cell type enrichments for M3 and M8. (**K**) RBP target enrichment for these modules include targets of *RBFOX2*, *CELF2* and *ELAVL2* (an nELAVL) which are hub isoforms in M3 and M8. (**L**) Transcript models for *ELAVL1* hub isoforms – arrow highlights the primate/human-conserved sequence missing from these 3’UTRs.

In contrast to geneExpr and isoExpr networks, the isoUsage network did not display strong cell type-specific enrichments (**Fig. 4A** and **fig. S6**), but instead was better defined by RBP isoform usage patterns and showed higher enrichment for RBP targets (**Fig. 4B** and **fig. S6B**). A detailed examination of isoUsage modules revealed expected enrichment patterns with modules exhibiting GZ/CP specificity enriched for targets of established progenitor- or neuronal-enriched RBPs such as *PTBP1*, *SRRM4*, *PTBP2,* and *RBFOX1/2/3* (**Fig. 4C**). However, many modules were also significantly enriched for targets of RBPs less studied in brain development, such as *SAFB*, *UTP18* and *SRSF9* (one-sided Fisher’s exact test, FDR-corrected P<0.05; (*50*)). Below, we focus on two module pairs with RBP isoforms in the top 30 hub transcripts and concomitant enrichment for their targets, and highlight example genes for which DTU informs neurodevelopmental processes.

IsoUsage.M1/M2 showed reciprocal GZ/CP specificity and concordant pathway enrichments for progenitor cell function and neuronal morphogenesis pathways, respectively (**Fig. 4C-E**). We also observed enrichment for neuronal markers in isoUsage.M1, reflecting the presence of neurons in the intermediate zone separating GZ and CP. Across isoUsage.M1/M2, we identified two novel NIC hub isoforms of the RBP *ELAVL2* (**Fig. 4C**) and concomitant enrichment of their targets (**Fig. 4F**). Inclusion of exon 2 in the CP-enriched isoUsage.M2 isoform (TALONT000708623) alters the translational start, adding 29 amino acids to the RNA-recognition motif (RRM) and potentially altering *ELAVL2’s* RNA metabolism function. Additionally, we found that in both modules, the *ELAVL2* hub isoforms contained an alternative 5’ TSS (**fig. S7F**) which may serve a regulatory function, as recent work demonstrated translational regulation of another ELAVL family member, *ELAVL4*, at its alternative 5’ UTRs (*62*). Consistent with GZ/CP DTU of BAF complex proteins (**Fig. 3F**), we identified three hub isoforms of *SMARCE1* (**Fig. 4G**), differing in the first four exons encoding an intrinsically disordered region (IDR). The isoUsage.M1 isoform encodes the full-length protein, whereas the two isoUsage.M2 isoforms lack either all or part of the first IDR, a low complexity protein domain (*63*) which mediates binding with higher-order complexes involved in chromatin remodeling and RNA splicing (*64*, *65*). Loss of all or part of the IDR in CP-enriched isoUsage.M2 suggests that *SMARCE1* may change its interaction with the BAF complex and/or associate with other protein complexes during neurogenesis.

In a second module pair, isoUsage.M3/M8, we observed complementary GZ/CP specificity and pathway enrichments, (**Fig. 4H**) and identified multiple RBP hub transcripts (*ELAVL1*, *RBFOX2*, *CELF2*) (**Fig. 4I-J**) with concomitant enrichment of their targets (**Fig. 4K**). While isoUsage.M3 was enriched in mRNA metabolism and RNA splicing processes, isoUsage.M8 was enriched in cytoskeletal function and cell projection organization (**Fig. 4I**), processes important in neuronal migration and maturation (*66*, *67*) and which match its CP enrichment (**Fig. 4J**). The M3 *ELAVL1* isoform encodes a short transcript containing only one of three RRMs, potentially affecting protein function (**Fig. 4L**). *ELAVL1* is also known to bind the 3’UTRs of target transcripts, including its own, to increase their stability (*68*) Both M3/M8 *ELAVL1* isoforms contain a primate/human-conserved novel 3’UTR with a ∼200bp intron spliced out, potentially affecting auto-regulation (**Fig. 4L** and **fig. S7G**). Altogether, these networks refine our understanding of the specific “RNA regulons” active in the developing brain (*46*).

### Isoform Expression at Single-Cell Resolution

Specific patterns of gene expression shape the differentiation and function of neural cells. Although gene expression in the developing neocortex has been extensively profiled at the single-cell level, isoform expression has yet to be systematically characterized. To gain single-cell resolution, we leveraged the recently developed ScISOrSeq (*20*) approach to profile >7,000 single cells across an additional 3 unique donor samples derived from microdissected GZ and CP regions (**Fig. 1A**; **fig. S8**). Barcoded full-length single-cell cDNA libraries generated using Drop-seq, with incorporated UMIs as published (*3*), were used as input to generate >26.4M high-quality PacBio CCS reads. To obtain cell-type specificity, cell-barcodes were matched to the high-depth, short-read sequencing dataset previously published on the same libraries (*3*). Of 7,189 individual single cell full-length transcriptomes, 4,281 had matching barcodes from short-read sequencing (**figs. S8 and S9A; table S5A**). All subsequent analyses were performed on this matched subset of the scIso-Seq data.

Following strict quality control and downstream processing (*36*) (**figs. S8** and **S9A-B**), we detected on average 530 unique transcripts per cell, mapping in aggregate to 18,541 genes and 138,497 unique isoforms (**fig. S9C, data S2**). We observed high concordance between pseudo-bulk short-read and long-read based gene expression (R = 0.9, p<2.2e^−16^) and detection (R = 0.92, p<2.2e^−16-^) (**fig. S9D** and **E**), and high inter-donor reproducibility (R = 0.84-0.87, p<2.2e^−16-^) (**fig. S9F**), demonstrating the robustness of the data. Similar to the bulk tissue transcriptome, the majority of detected isoforms (71.7%) were previously unannotated (**fig. S9G**). We found broad support for these isoforms in bulk tissue IsoSeq and in independent long-read datasets, with >80% matching both 5’ and 3’ end termini (**fig. S9H**), >75% containing CAGE or ATAC-seq peaks near the TSS, >85% supported by nearby polyA sites or motifs (**fig. S9I**), and ∼87% of splice junctions detected in bulk IsoSeq (**fig. S9J**). Altogether, 67,183 isoforms detected by scIso-Seq (49% of total) fully match isoforms detected in bulk tissue and 40% match independent datasets (**fig. S9K**), including those containing 2,452 out of 5,165 previously unannotated spliced-in exons (**table S5B**), a handful of which were also validated by RT-PCR (**fig. S9L**).

We next connected single cells with their specific cellular identities. Previous unsupervised graph-based clustering in Seurat (*69*) using high-depth short-read RNA-Seq identified 16 transcriptionally distinct cell-type clusters in the developing human neocortex (**Fig. 1A**) (*3*). Through barcode matching, we detected cells from all 16 clusters (**Fig. 5A**), allowing us to construct cell-type-specific isoform expression profiles (**table S5** and **Fig. 5B**). Comparing isoform expression diversity across cell types, we observed that excitatory neuron clusters, in particular those corresponding to newly-born migrating (ExN) and maturing neurons (ExM), harbored the largest number of isoforms (**Fig. 5C**). This was not due to differences in sequencing depth across clusters (**fig. S9M**) or gene detection (**fig. S10A**). These same cell types exhibited the greatest diversity of unannotated expressed isoforms (**Fig. 5C**), highlighting a role for these transcripts in early neuronal maturation processes.

**Fig. 5.**
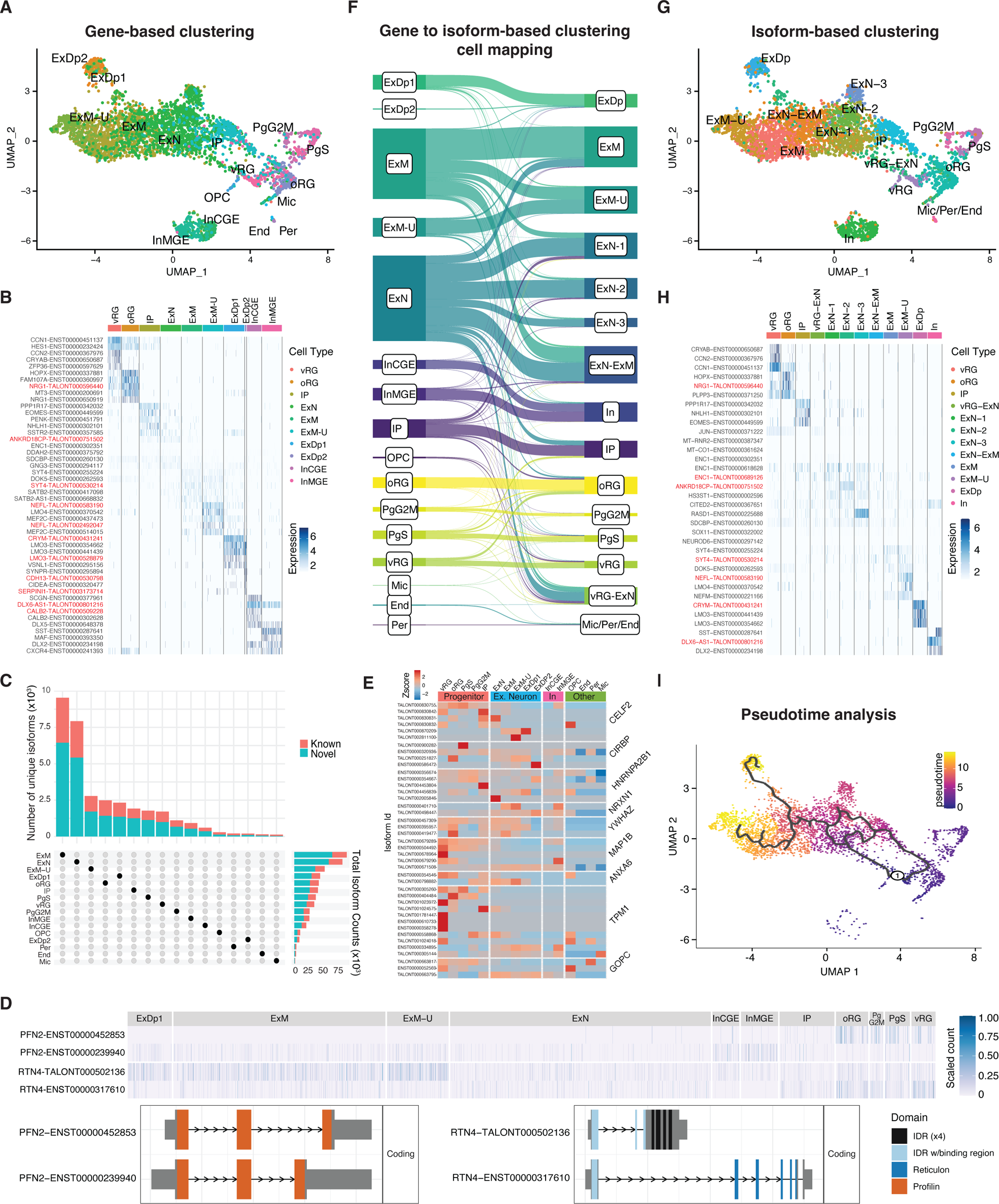
Cell type-specific isoform diversity in the developing human cortex. (**A**) Uniform manifold approximation and projection (UMAP) plot of 4,281 cells detected by both 3’ end short-read sequencing and by scIso-Seq. Each dot represents a single cell, colored by its corresponding cluster. UMAP position of the cells is calculated based on isoform expression while cluster labels are as previously defined ((*3*)). vRG = ventricular radial glia, oRG = Outer Radial Glia, PgG2M = Cycling progenitors (G2/M phase), PgS = Cycling progenitors (S phase), IP = Intermediate Progenitors, ExN = Migrating excitatory, ExM = Maturing excitatory, ExM-U = Maturing excitatory upper enriched, ExDp1 = Excitatory deep layer 1, ExDp2 = Excitatory deep layer 2, InMGE = Interneuron MGE, InCGE = Interneuron CGE, OPC = Oligodendrocytes Precursor Cells, End = Endothelial, Per = Pericyte, Mic = Microglia. (**B**) Heatmap showing differentially expressed isoforms across cell types defined by gene-based clustering. Novel isoforms are shown in red. (**C**) Distribution of isoforms across cell types shows greater diversity of isoforms in newborn migrating (ExN) and maturing excitatory neurons (ExM) compared to other cell types in mid-gestation human cortex. (**D**) Isoforms of PFN2 and RTN4 differentially expressed across cell types along with their predicted functional consequences. Isoform ENST00000239940, predominantly expressed in neurons, is predicted to encode IDR protein domains not found in the progenitor-enriched ENST00000452853 isoform. The novel isoform TALONT000502136 is enriched in neurons, whereas the progenitor enriched ENST00000317610 isoform is longer and contains multiple reticulon protein domains. (**E**) Heatmap showing a subset of isoforms with differential usage across cell types (DTU). (**F**) River plot showing mapping of cells from gene-based (left) to isoform-based (right) clustering. Each line represents a single cell. (**G**) UMAP of cells clustered based on isoform expression as measured by scIsoSeq. Additional stages of excitatory neuron maturation can be defined using isoform-level data. (**H**) Heatmap showing differentially expressed isoforms across cell types defined by isoform-based clustering. Novel isoforms are shown in red. (**I**) Cell lineage trajectory analysis (Monocle3) shows direct neurogenesis through vRG-ExN cells and indirect path through IP cells.

Selective isoform expression across different cell types has been reported in the adult brain for a few genes (*19*, *70*, *71*) with potential implications for neuropsychiatric disorders (*25*). Thus, we next sought to systematically characterize isoform expression and utilization across cells in the developing neocortex. We first characterized patterns of DTE between each cell-type cluster versus all other clusters (*36*) (**table S5C**) to identify transcripts that define cellular identities and can serve as molecular markers (**Fig. 5B**). As expected, the majority of these transcripts belonged to genes previously identified as canonical markers of the respective cell-types, including *HES1* (RG), *CRYAB* (vRG), *HOPX* (oRG), *EOMES* (IP), *LMO3* (ExDp), and *SATB2* (ExM-U), among others (*3*). Of 1,040 transcripts enriched in specific cell-types, 257 (24.7%) corresponded to isoforms newly identified in this study (**table S5C**). Among the top enriched transcripts for each cluster, we identify novel isoforms of *NRG1*, *LMO3*, *NEFL,* and *SYT4* enriched in oRG, ExDp, ExM-U, and ExM cell classes, respectively, all of which have established roles in brain development and function (**Fig. 5B, table S5C;** (*72–77*)).

We next conducted a pairwise DTE analysis to identify transcripts changing across specific cell-type transitions, detecting 409 transcripts corresponding to 147 genes (**table S5D,** P_nominal_<0.05). Focusing on genes with multiple isoforms showing dynamic expression between progenitors and neurons, we observed isoforms of *PFN2*, which functions in actin polymerization dynamics and morphogenesis (*78*), with opposing expression patterns between progenitors and neurons (**Fig. 5D**). Similarly, we identified progenitor and neuron-specific isoforms of *RTN4*, a canonical regulator of axon growth and neuronal migration (*79*, *80*) (**Fig. 5D**). Together, these examples highlight changes in isoform expression across cell-types and developmentally-relevant transitions with putative consequences to the structure or stability of their encoded protein products.

Given the degree of isoform switching observed between GZ and CP (**Fig. 3**), we sought to quantify similar events across individual cell types (*36*). We identified 1,695 genes where the proportion of expressed isoforms for a given gene differed across at least two cell types (single-cell DTU; **Fig. 5E** and **table S5E**). These instances represent switches in isoform utilization across cell types that may be missed by traditional DGE analyses. Of the 2,284 specific transcripts exhibiting DTU across these 1695 genes, 48.5% showed proportional differences in progenitors and 43% in neurons, with an average number of 221 DTU transcripts per cell-type with roughly similar distribution across these cell classes. DTU genes were enriched in regulation of mRNA splicing (*CELF2*, *CIRBP*, *HNRNPA2B1*), cell division, regulation of synapse maturation (*NRXN1*, *YWHAZ*), and GO categories related to cytoskeleton dynamics and vesicle transport (*MAP1B*, *ANXA6*, *TPM1*, *GOPC*) (**Fig. 5E** and **fig. S10B**). Consistent with these results and a role for isoform switching in cell identity, GZ/CP DTU transcripts primarily clustered by expression across progenitors, neurons, or support cells (**fig. S10C**).

### Additional Cell Types Uncovered from Isoform-level Clustering

Given the broad changes in isoform diversity and expression observed across cells, we leveraged these data to expand current cell-type classification catalogs. Re-clustering cells based on isoform expression yielded 15 highly-stable clusters largely mapping to many of the same cell classes as defined by gene-based clustering (*36*) (**Fig. 5 F-H** and **fig. S10D**). However, progenitors transitioning into neurons and early-born excitatory neurons were split into additional clusters, providing higher-resolution cell maturation stages than those observed by traditional gene-based clustering. In particular, newborn migrating neurons (ExN) split into three clusters (ExN1-3), encompassing cells previously annotated to IP, ExN, and ExM clusters, and two additional new clusters, vRG-ExN and ExN-ExM, representing cells in states on either side of a maturity spectrum centered around ExN cells (**Fig. 5 F-G**). This was supported by pseudotime lineage inference analysis, whereby vRG-ExN cells represented a path of direct neurogenesis from vRG cells distinct from another path through oRG cells, and where ExN-ExM cells preceded the most mature neuronal clusters (**Fig. 5I**). To better understand the molecular programs and markers of these new cell states, we repeated DTE analyses (**table S5C**). Across ExN clusters, isoforms of *ENC1, ANKRD18CP,* and *RASD1* were enriched in ExN1, ExN2, and ExN3 cells, respectively (**fig. S10E-F**). Moreover, vRG-ExN cells were defined by a majority of transcripts involved in mitochondrial metabolism, consistent with a recent report of the role of mitochondria in regulating neuronal maturation (*81*). Overall, the increased resolution in ExN and ExM cells obtained from isoform-based clustering matches the observed increase in isoform diversification in those cells (**Fig. 5C**) and supports a role for this mechanism in the early processes of neurogenesis.

### Isoform-Centric Localization of Convergent Risk Gene Mechanisms

We next performed enrichment analyses to localize rare-variant association signals from large-scale whole exome and genome sequencing studies of neurodevelopmental and psychiatric disorders, including ASD (*82*), NDD (*82*, *83*), SCZ (*84*), bipolar disorder (BIP) (*85*), and epilepsy (*86*). Risk genes for NDD, ASD, and DDD (Developmental Disabilities) had significantly more isoforms (log2 scale; OR’s 1.14-1.4; q’s<3e-4) and exons (log2 scale; ORs 1.19-1.64; q’s<0.01) compared to non-disease genes (**Fig. 6A**, **fig. S11A, table S6A;** FDR-corrected p-values from logistic regression, correcting for gene length and coding length). These associations were not observed for epilepsy (ORs 1.14, 1.22; NS), BIP (ORs 1.12, 1.22; NS), or SCZ (ORs 1.4, 1.56; NS). Disease-associated genes showed significant overlap with those exhibiting DGE, DTE, and DTU during cortical neurogenesis (logistic regression, FDR-corrected P<0.05; **Fig. 6A** and **fig. S11B**). Overall, these associations were observed mainly for genes and isoforms upregulated in the CP (DGE.up, DTE.up), indicative of neuronal expression, and were shared across NDD, non-syndromic ASD, DDD, and epilepsy, but not other diseases. In particular, NDD genes were enriched for those exhibiting DTE or DTU, but not changing in overall gene expression (DTU.not.DGE, DTE.not.DGE). Consistently, at the single-cell level we observed that NDD genes were significantly enriched in DTU (logistic regression, FDR-corrected P<0.05; **Fig. 6B**, “SingleCell-DTU”), and many NDD and ASD genes showed DTU across cell types in the developing neocortex (**Fig. 6C; fig. S12A**). NDD, DDD, and ASD gene isoforms were primarily enriched in excitatory neurons (ExM-U, ExDp, or ExM) (**Fig. 6B; fig. S12B**). However, isoforms of NDD genes and those causing syndromic forms of ASD were also enriched in mitotic progenitors and radial glia (**Fig. 6B; fig S12B**), as expected from the broader phenotypic spectrum in these disorders. Finally, NDD genes were enriched in ExN1, the newly-defined cell state based on isoform-level quantifications (**Fig. 6B** and **Fig. 5**).

**Fig. 6.**
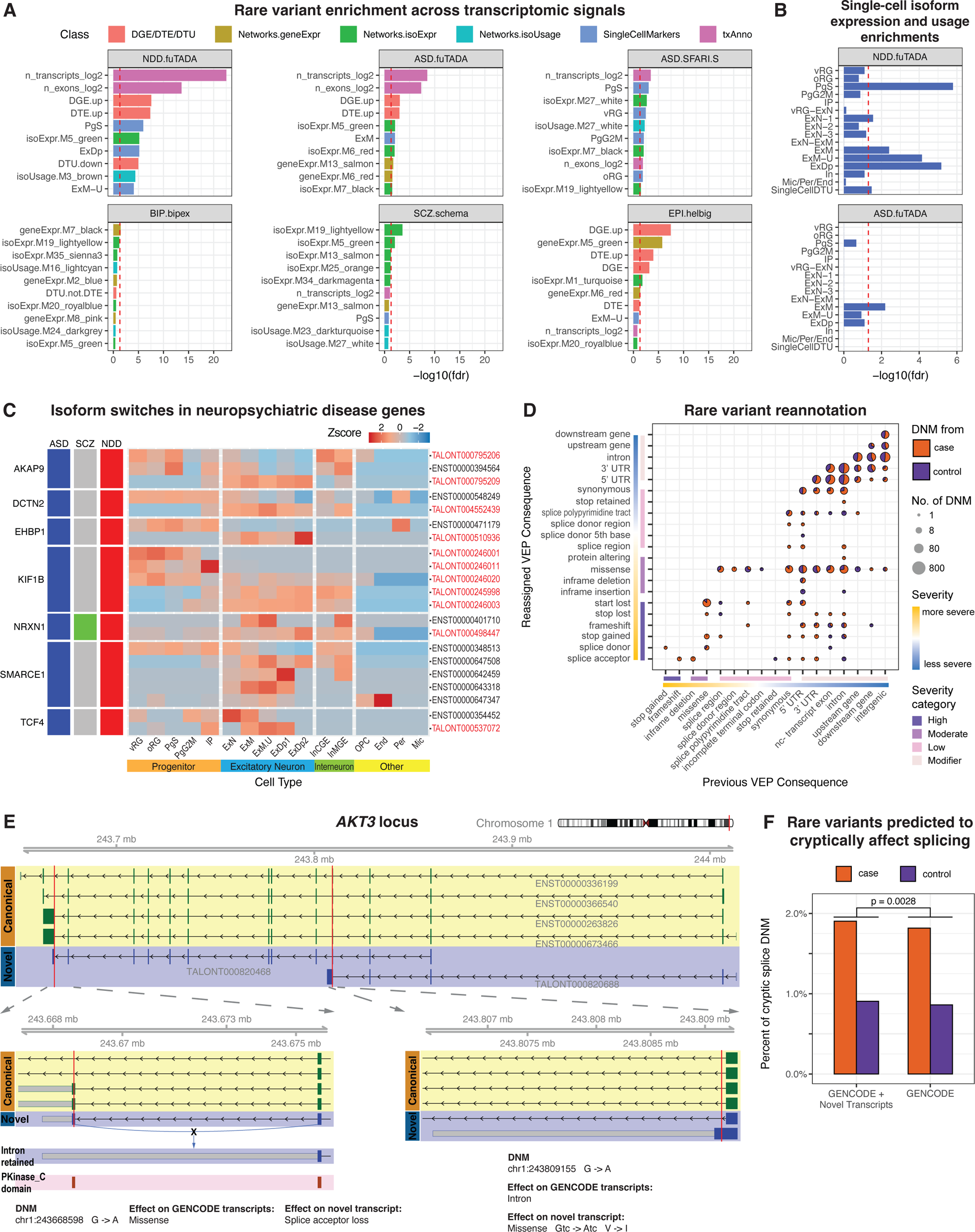
Isoform-centric contextualization of neurogenetic risk mechanisms. (**A**) Enrichment of transcriptomic features, differential expression analyses across cortical regions and cell types or isoform expression and usage networks with neuropsychiatric disorders. Red line indicates the FDR-corrected significance threshold. (**B**) Cell type enrichments indicate differential isoform expression and utilization in NDD and ASD. (**C**) Heatmap of isoforms from several NDD and ASD risk genes showing differential usage across the cell types of the developing cortex. Novel isoforms are labeled in red. (**D**) Number of variants that were reassigned to a more severe consequence after taking into account newly identified isoforms in this study. The size of the dots represents the number of variants in each category. The color of the dots indicates the source of the variants, i.e. DNM from case or control. Colored bars along the axes indicate both the severity of the consequences on a continuous scale, and the severity categories on a discrete scale, as defined by VEP. Reassignment to a different severity category may be more impactful than reassignment within the same category. (**E**) The *AKT3* gene locus with representative canonical isoforms and two novel isoforms identified from this study. Red vertical lines indicate the position of case DNMs that affect this locus. The affected regions are highlighted in the lower panels. Lower right panel, a DNM falls in the intron region in canonical protein isoforms, while leads to missense mutation in a novel protein isoform. Lower left, a DNM causes the loss of nearby splice acceptor and intron retention only in a novel protein isoform. The retained intron leads to shortened coding sequence and eliminates part of the protein kinase C-terminal domain. (**F**) Proportion of de novo mutations predicted to cryptically affect splicing, with or without the annotation of newly identified isoforms from this study.

We next leveraged our gene/isoform co-variation networks to localize disease gene convergence at the molecular level. Disease gene signal mainly coalesced among isoExpr modules (51.9% at p_nominal_<0.05), followed by isoUsage (26.8%) and geneExpr (21.3%) modules (**fig. S13**). NDD, DDD and non-syndromic ASD shared overlapping molecular signatures with modules mainly enriched for neuronal markers (**fig. S13**) and isoExpr modules regulating chromatin/histone modification (isoExpr.M7; **Fig. 6A and fig. S7B**) and RNA metabolism/splicing (isoExpr.M11/ M10/M28; **table S6A and fig. S7C-E**). Notably, isoExpr.M11 contained a hub isoform of the ASD-associated RNA helicase *DDX3X* with concomitant enrichment for its targets (**fig. S7E**). NDD and ASD genes were also enriched across excitatory neuronal modules regulating cytoskeleton, synaptic vesicles, and neurite morphogenesis (isoExpr.M30/M24, **table S6A** and **fig. S7D**) and ribosomal RNA processing and chromatin (isoUsage.M29, **table S6A and fig. S7C**). These findings support a major role for isoform expression and diversification in neuropsychiatric disease mechanisms during development, consistent with recent findings (*25*), regulating chromatin remodeling, cytoskeletal dynamics and RNA processing.

### Reprioritization of De Novo Variants in Individuals with NDDs

Finally, to move from population to individual genetic risk mechanisms, we used our atlas to reinterpret de novo, noncoding genetic variants identified in large-scale sequencing studies of ASD (*87–89*) and intellectual disability/developmental disorders (ID/DD) (*89*). We reasoned that some variants previously disregarded as noncoding may actually fall within the >27MB with newfound transcriptional activity in our data or disrupt newly identified splice junctions. To test this hypothesis, we complemented the Gencode v33 annotation with our newly identified protein-coding isoforms and re-annotated the set of compiled genetic variants (N_total_=272,187; N_case_=145,880; N_control_=126,307) using VEP. Altogether, this new annotation framework uncovered more severe consequences for 1.24% of all variants (**Fig. 6D**, **table S6B**). For example, we observed a novel *AKT3* isoform (TALONT000820688) with an alternative last exon extending the coding sequence (**Fig. 6E; right**) and expressed at comparable levels as known *AKT3* isoforms (**fig. S14**). This extended coding region overlapped a de novo mutation (DNM) from the ASD cohort (*90*), previously classified as a benign intronic variant but now predicted to cause a missense mutation in the novel AA sequence. In another example, a reported DNM from the ID/DD cohort (*91*) was predicted to cause a missense mutation in the *KLC1* protein based on known isoforms from Gencode (v33). The *KLC1* gene encodes a member of the kinesin light chain family, involved in microtubule cargo transport. We observed a *KLC1* isoform (TALONT000423578) with a novel TSS, supported by overlapping CAGE peaks (**fig. S15A**). This isoform was predicted to code for a protein with a novel start codon. Given the structure of this novel isoform, the DNM would lead to the loss of the start codon--a potentially more severe consequence. The protein encoded by TALONT000423578 had an alternative carboxy termini not observed in Gencode (v33) and strongly supported by our proteomics data (**fig. S15B**).

Cryptic splicing variation, in which splice-disrupting variants fall outside of the essential GT and AG dinucleotide motif, is a major recently uncovered mechanism underlying genetic risk for neurodevelopmental disorders, as exemplified by SpliceAI (*23*), a state-of-the-art deep neural network trained using pre-mRNA sequence to predict cryptic splice mutations. To determine whether our isoform-centric transcriptome could improve prediction of NDD-associated cryptic splice variants, we retrained SpliceAI with these annotations and overlapped its predictions with the compiled set of DNMs. With the addition of novel transcripts, a significantly larger proportion of variants were predicted to be cryptic splice variants (**Fig. 6F**, **table S6C** 1.44% with Gencode and novel transcripts, 1.37% with only Gencode transcripts, p = 0.0028, binomial test). By way of example, an ID/DD associated DNM was predicted to alter the splicing of a novel *AKT3* isoform (TALONT000820468), while it had no effect on the splicing of known *AKT3* isoforms (**Fig. 6E**). TALONT000820468 is the most highly expressed *AKT3* isoform detected (**fig. S14**). This variant caused the loss of the nearby splice acceptor and the retention of the last intron. The retained intron leads to shortened coding sequence and eliminates part of the protein kinase C-terminal domain (**Fig. 6E**). *AKT3* is a key regulator of the PI3K-AKT-mTOR pathway in the nervous system (*92*), and dysregulation of *AKT3* is associated with NDDs (*93*). Our findings highlight multiple de novo mutations may contribute to NDDs by affecting specific *AKT3* isoforms. More broadly, these results demonstrate that a more complete catalog of mid-gestation-brain-expressed full-length isoforms provides more granular molecular insight into the genetic risk mechanisms underlying neurodevelopmental disorders.

## Discussion

Here, we have provided a detailed view of the full-length, alternatively-spliced transcriptome in the developing human neocortex at midgestation, with regional and cell-type specificity. Although splicing and isoform regulation are known to be critical for proper neural development (*15*), and are strongly implicated in NDD risk (*21*, *23*, *25*), technical challenges have made it difficult to delineate the path from genetic mutation to functional isoform changes, in part due to reliance on short-read sequencing as well as incomplete genomic annotations. Using high-depth long-read sequencing, we identified 149,510 previously unannotated transcript isoforms in the developing human brain, extending by >27MB the transcriptionally active content of the genome and expanding the proteomic diversity of the human brain. The majority of these unannotated transcripts were validated across independent datasets and data modalities; the remaining fraction may be unique to the specific donors, developmental periods, and/or regions profiled here or, alternatively, represent false positives inherent to challenges of isoform annotation. Examining the functional consequences of novel isoforms and profiling their expression across a wider range of time periods, cell-types, and distinct genetic backgrounds, facilitated by decreasing sequencing costs, will be critical next steps for future studies to further refine these results. However, our analyses here provide considerable support for their functional importance during neurogenesis and in neurodevelopmental disorders.

Assessing differential transcript-isoform activity across the developing cortex, we found wide-ranging changes in isoform expression and usage implicating chromatin remodeling via the BAF complex, and cytoskeletal dynamics important for neuronal morphogenesis. Isoform switching during corticogenesis implicated known neuronal splicing regulators, as well as RBPs previously not studied in context of brain development, including KHSRP, SUPV3L1, and SRSF9. APA analysis of GZ/CP differential DTU genes supported previous work showing that 3’UTR lengthening in differentiated cells is conserved throughout many cell lineages including neurons (*94*, *95*). Though there are many individual examples of APA in neuronal RBP genes (*94*, *96*, *97*), our analysis found that APA of RBP genes occurs on a large scale during human corticogenesis, indicating that tight regulation of RBP activity is important for neurogenesis.

In network analyses, we found that while geneExpr and isoExpr networks are cell-type driven, the isoUsage network was defined by RBP regulatory dynamics. IsoUsage modules contained hub isoforms with putative structural and regulatory differences including those predicted to alter protein domains of the NDD risk gene *SMARCE1*, and encoding primate and human-specific novel 3’UTR sequence of the RBP *ELAVL1.* While many developmental processes are conserved across vertebrates in early neural development, primate- and human-specific differences are important to understand given the unique cell types found in the expanded cortices of these species (*98*, *99*). Correspondingly, isoform-level single-cell transcriptomics demonstrated differential isoform expression and usage across cell types and enabled the identification of additional cell states in newborn excitatory neurons (ExN 1-3) as well as states encompassing the transition from progenitor to neuron and neuronal maturation (vRG-ExN and ExN-ExM). Together, these results increased the catalog of isoforms expressed during corticogenesis and strongly implicated splicing and RBP regulation of isoform expression and usage in neurogenesis.

The data generated in this study can inform current and future genome-wide disease risk association studies. We show that genes associated with NDD, ASD, and DDD exhibit increased isoform diversity, and NDD and ASD rare variants are enriched for isoform expression and usage changes during corticogenesis. Finally, we used our isoform-centric atlas to re-annotate and re-prioritize thousands of de novo ASD/IDD rare variants. The large number of previously unannotated transcripts identified here suggests that the functional consequences of many variants may have been missed using previous incomplete annotations. In sum, our results have broad implications for understanding cell fate specification in the developing human brain and for comprehensive interpretation of the genetic risk mechanisms underlying developmental brain disorders.

## Materials and methods summary

Detailed materials and methods can be found in the supplementary materials. Human mid-gestation cortical tissue samples were obtained from the UCLA Gene and Cell Therapy Core in accordance with the IRB and the UCLA Office of Human Research Protection regulations, with full informed consent from the parent donors. GZ and CP regions of 15-17 PCW cortices were microdissected and processed for bulk IsoSeq or scIsoSeq. For bulk IsoSeq, PacBio Sequel IIe platform was used to generate ∼38.5 million high-quality, full-length reads, which were filtered, aligned to the human reference genome, and analyzed using TALON, TranscriptClean, and Gencode v33 to identify and quantify known and novel genes/isoforms. Differential gene expression, transcript usage, and alternative polyadenylation site usage analyses were conducted using DESeq2, IsoformSwitchAnalyzeR, and DaPars2, respectively. Pathway enrichment was performed using gProfileR, while RBP target gene sets were analyzed for overlap with gene lists or module-associated genes/isoforms from bulk isoform sequencing or network analyses using Fisher’s exact test and FDR-correction. TransDecoder, CPAT, and Comet were used to predict ORFs, assess coding potential, and validate novel proteins, respectively. WGCNA was used to generate gene and isoform expression, and isoform utilization modules, followed by over-representation and GO analysis. scIsoSeq libraries were prepared from captured single-cell full-length cDNA (*3*), and sequenced on Sequel I/II platforms. Reads were mapped to the human genome reference using minimap2 and isoforms were called with TALON. Seurat was used for clustering and differential expression analysis and Monocle3 for lineage trajectory analysis. Single-cell differential transcript usage analysis was conducted using binomial regression and p-value correction. RT-PCR was used to validate novel exons. Enrichment analyses were performed to associate rare-variant signals from neuropsychiatric disorders with transcriptome features, gene/isoform modules, or differentially expressed genes/isoforms using logistic regression controlling for gene and coding length, and transcript expression. ASD/IDD de novo variants(*87–89*) were annotated using Ensembl VEP or SpliceAI, with two rounds of annotation involving GENCODE v.33 GTF file and predicted ORFs from newly identified transcripts.

## Acknowledgments

We thank members of the Gandal, de la Torre-Ubieta, Li and Pasaniuc laboratories for helpful discussions and critical reading of the manuscript. We thank Dr. Geschwind for kindly providing bulk RNA and single-cell cDNA libraries used as input for long-read sequencing library preparation. Pac-Bio bulk tissue long-read library generation and sequencing were performed at the UC Davis Genomics Core and brain tissue was obtained from the UCLA CFAR (5P30 AI028697).

## Author Contributions

A.P., M.J.G and L.T.U. conceived and designed the study. C.V. and L.T.U. collected and processed the tissue specimens and dissected the samples for processing. A.P. and L.T.U. generated the library for single cell IsoSeq. A.P. processed the single-cell raw data and performed the validation experiments. C.J. and A.P. analyzed bulk tissue IsoSeq data, C.J and M.J.G. analyzed bulk tissue isoform switching, P.Z., X.W. and C.L. performed the proteomic analysis, N.G. performed the alternative polyadenylation analysis, C.V. performed the RBP associated analysis, C.V., L.T.U. and M.J.G. conducted network analyses, A.P., C.J., M.M. performed single cell IsoSeq analysis, M.K., D.V., A.P. performed single cell isoform switch analysis. X.G., J.J.L. perform single cell DTU analysis, A.P and L.T.U analyzed the DTU results, M.J.G, L.T.U., P.Z., K.H., C.J., A.P., B.P. performed the disease enrichment analysis. A.P., C.V., M.J.G., L.T.U. interpreted the data; A.P., C.V., P.Z., M.J.G., L.T.U. wrote the manuscript.

## Data and materials availability

Controlled-access bulk and single-cell IsoSeq data is available at https://doi.org/10.7303/syn4921369 and https://assets.nemoarchive.org/dat-rhocguc to investigators subject to approval by the NIMH Repository and Genomics Resources (NRGR).

A UCSC track hub browser containing mid-gestation neocortex isoforms is available at: https://genome.ucsc.edu/cgi-bin/hgTracks?hubUrl=https://raw.githubusercontent.com/ashokpatowary/Dev_Brain_IsoSeq/main/hub.txt&genome=hg19&position=lastDbPos

## Code availability

Code used to process the data and generate all figures in this manuscript is available at https://github.com/gandallab/Dev_Brain_IsoSeq and from Zenodo at https://zenodo.org/badge/latestdoi/614018591

## Supplementary Materials

Materials and Methods

Figs. S1 to S15

Tables S1 to S8

Data S1, S2

References 100 - 119

---

**Summary:**
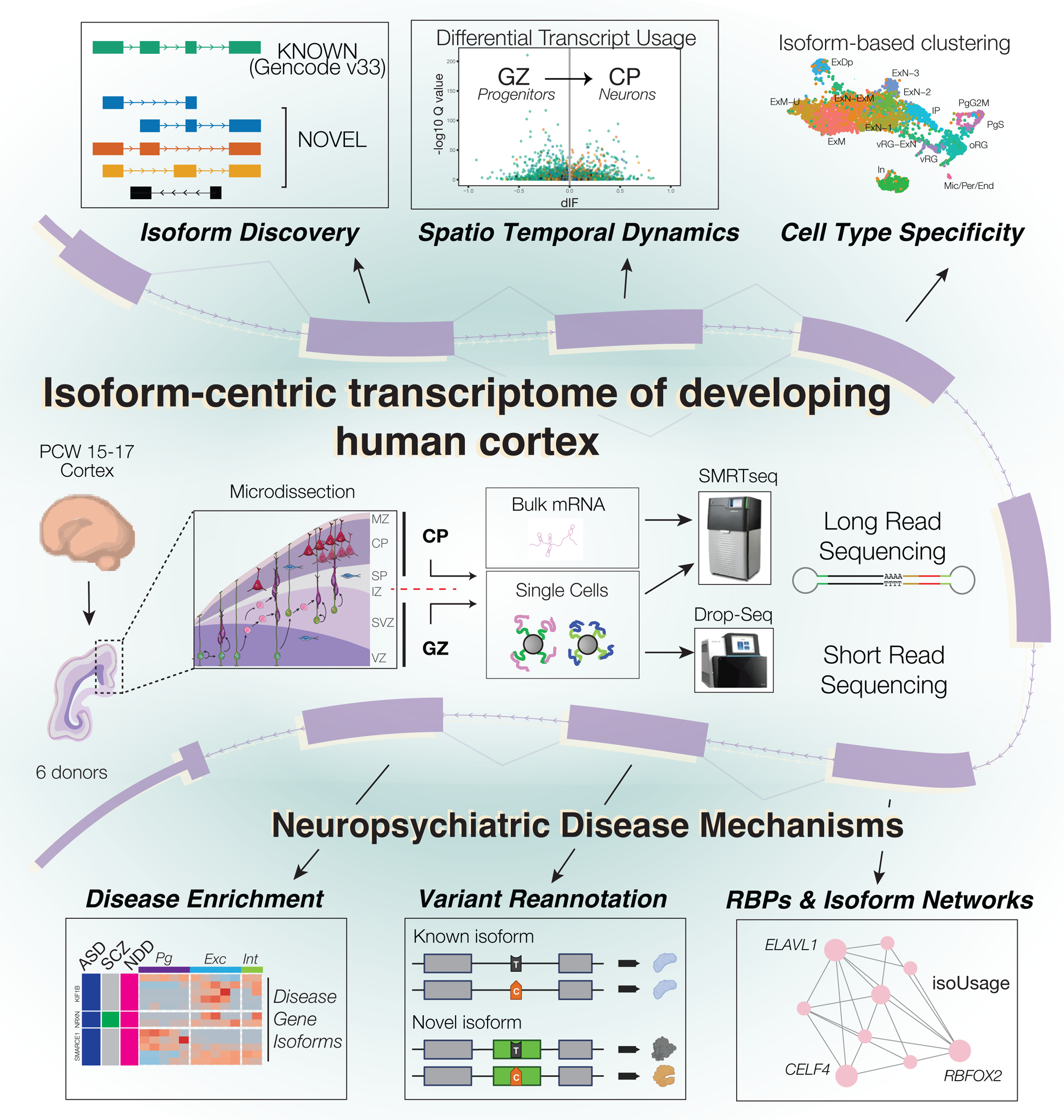
An isoform-centric transcriptome of the developing human neocortex informs mechanisms of neuropsychiatric disease. We provide a systematic characterization of transcript-isoform diversity in the developing human neocortex at tissue and single-cell resolution using PacBio IsoSeq. We identify thousands of novel isoforms with specific regional (GZ/CP) and cell type expression coalescing into networks driven by cell type identity and RNA binding protein regulation. This resource reveals substantial contributions of isoform switching to cellular identity and elucidates novel genetic risk mechanisms for neurodevelopmental and neuropsychiatric disorders, including a re-annotation of thousands of de novo rare variants with potential clinical implications.

