## Supplementary material for "Developmental isoform diversity in the human neocortex informs neuropsychiatric risk mechanisms"

**The PDF file includes:**

Materials and Methods

Figs. S1 to S15

Tables S1 to S8

References 100 - 119

**Other Supplementary Materials for this manuscript include the following:**

Data S1, S2

### Materials and Methods

#### Tissue acquisition, dissection, and sample processing

De-identified human mid-gestation brain tissue for all studies was obtained from the UCLA Gene and Cell Therapy Core according to IRB guidelines and to the legal and institutional ethical regulations of the UCLA Office of Human Research Protection. Full informed consent was obtained from all of the parent donors. No known major pathogenic CNVs implicated in neuropsychiatric disorders were found in any donors.

For bulk Iso-Seq, previously-captured RNA (*37*) from 3 donors aged 15-17 PCW was generated as described before. Briefly, for each donor, a 1-2mm slice of cortex was prepared and then microdissected along the intermediate zone with a razor blade under a dissection microscope to obtain two regions, the progenitor-enriched germinal zone [GZ; encompassing the ventricular zone (VZ), subventricular zone (SVZ), and part of the intermediate zone (IZ)] and the postmitotic neuron-enriched cortical plate (CP; encompassing part of the IZ, the subplate, cortical plate and marginal zone). RNA was extracted independently from each region of the donor samples using the QIAGEN miRNeasy Mini Kit. RNA concentration was measured using a Qubit 4.0 fluorometer (Thermo Fisher Scientific), and sample RNA quality was evaluated by RNA Integrity Number (RIN) on an Agilent tapestation system. All RNA samples were of high quality, with RIN values > 8.0.

For single-cell Iso-Seq, previously-captured(*3*) single-cell full-length, barcoded cDNA libraries from 3 donors aged 15-16 PCW were used, prepared as described before. Briefly, cortical tissue was microdissected as above. Dissected GZ/CP regions were treated with papain (Worthington) for enzymatic dissociation and filtered into a pure homogeneous cell suspension through a 40um strainer and an ovomucoid gradient. Cell survival (90%–95%) and yield were quantified with Trypan blue staining before proceeding with cell capture. Drop-seq v3.1 was used to capture and barcode individual whole cells and to generate full-length cDNA libraries with unique molecular identifiers (UMI) for each RNA molecule.

#### IsoSeq library preparation, sequencing, read mapping and QC

From each of the bulk tissue samples, uniquely barcoded full-length cDNA libraries were synthesized using NEBNext Single Cell/Low Input Kit with 300ng of RNA as input. Briefly, RNA was incubated with NEBNext Single Cell RT Primer Mix for 5 min at 70°C followed by addition of Single Cell RT buffer and Single Cell RT enzyme mix and incubation for 75 min at 42°C. Afterwards, template switching oligo (TSO) was added, followed by 15 min incubation at 42°C. cDNA was purified using 1.0x ProNex beads. Subsequently, purified cDNA was further amplified for 12 cycles with a specific barcode-containing primer and the Iso-Seq Express cDNA PCR primer. Amplified cDNA was further purified by 1.0x ProNex beads at a 1:1 ratio.

Library generation and sequencing were performed at the UC Davis Genomics Core. For SMRTbell library preparation, pairs of samples were pooled in equimolar concentrations and libraries were prepared using the SMRTbell Express Template Prep Kit 2.0 following the manufacturer’s instructions, including damage repair, end repair, A-tailing and ligation of overhang adapter followed by purification of the libraries using ProNex beads. Library concentration was assessed by Qubit and size distribution was assessed by Agilent Bioanalyzer. Final SMRTbell Sequencing complexes were generated using Sequel II Binding Kit 2.0, Sequencing Primer v4.0, and Sequel II Polymerase 2.0, and then purified using ProNex beads. Pooled samples were then sequenced across several 8M SMRT-cell on the PacBio Sequel IIe platform with 20-hour movie collection. A total twelve 8M SMRT-cells were used to generate the IsoSeq data.

SMRT-Link (version 10.2.0) was used to call circular consensus sequencing (CCS) reads using default parameters (--all --minLength 10 --maxLength 50000 --minPasses 3 --minSnr 2.5 --minPredictedAccuracy 0.99). Across the twelve 8M SMRT cells sequenced, we generated ~38.5 million full-length, non-chimeric reads. As the libraries were generated with duplex samples, demultiplexing was performed to separate the reads from six samples across four SMRT cells. The lima package was used on the demultiplexed CCS reads to remove primers and adapters, and the isoseq3 tool to filter out reads without polyA sequence. Filtered reads were mapped to the human reference genome GRCh37 using minimap2 (version 2.24) with parameters “-ax splice:hq -uf --secondary=no --MD”. Finally, we generated a merged, filtered BAM, containing only reads with a predicted read accuracy of 0.99 or above (PacBio hifi).

#### IsoSeq transcript discovery and quantification

We used TALON (v5.0) and its associated utilities for identifying and quantifying known and novel genes/isoforms. We first ran TranscriptClean on the mapped reads using default parameters to correct mismatches, microindels, and noncanonical splice junctions based on the reference genome. To remove incomplete reads that may be caused by internal polyA priming caused by A-rich sequences, we used talon_label_reads to compute the fraction A values for a post-transcript interval of 20bp and removed reads with a fraction greater than 0.75. We then initialized a TALON database using Gencode v33 as our reference annotation. We fed our mapped reads into the database using the talon command with parameters “--cov 0.85 --identity 0.8”. TALON classified isoforms into multiple categories: known, ISM (incomplete splice match); NIC (novel in catalog); NNC (novel not in catalog); Genomic; Intergenic and Antisense. A GTF annotation file was generated using talon_create_GTF module for isoforms passing the following criteria:

(1) Isoforms labeled as known
(2) Isoforms labeled as novel (e.g., ISM/NIC/NNC/Genomic/Antisense/Intergenic), ***and***
    supported by 3 independent reads, ***and***
    (a) Observed in two independent donors, ***or***
    (b) Exhibiting independent 5’ and 3’ support, ***or***
    (c) Present in an independent long read isoform database.

For analysis of 5’ and 3’ support, we used SQANTI3 (*35*) on the unfiltered set of transcripts from the TALON database. 5’ support was based on CAGE peaks found in the refTSS v3.3 annotation (*100*), while 3’ support was determined based on either polyA site data from PolyASite 2.0 atlas (*41*) or nearby polyA motifs detected by SQANTI3. Independent long read isoforms came from PacBio’s publicly available Alzheimer’s Disease human brain ([https://downloads.pacbcloud.com/public/dataset/ Alzheimer2019_IsoSeq/](https://downloads.pacbcloud.com/public/dataset/Alzheimer2019_IsoSeq/)) or the Universal Human Reference RNA datasets (<https://github.com/PacificBiosciences/DevNet/wiki/Sequel-II-System-Data-Release:-Universal-Human-Reference-(UHR)-Iso-Seq>). The resulting whitelists were used to generate filtered abundance files for transcript quantification as well as custom-filtered GTF annotations for all subsequent analyses, including to identify putative novel transcribed exons. With this filtered transcriptome, we performed a final check for external support based on adult and fetal CAGE peaks (*101*) within +/- 200bp of the TSS (*40*, *100*), ATAC-Seq peaks (*37*–*39*) within +/- 500bp of the TSS and polyA sites (*41*) and polyA motif data from SQANTI3. Additionally, splice junction information was obtained by SQANTI3. Splice junctions were categorized into canonical and noncanonical based on the presence of specific di-nucleotide pairs at the boundaries of introns. Canonical splice sites are well-established and the most commonly observed sites, comprising GT at the donor site (immediately after the 5' end of the exon) and AG at the acceptor site (prior to the 3' end of the exon). Additionally, the less common variants GC/AG and AT/AC are also considered canonical, while all other dinucleotide combinations are classified as noncanonical. For the novel spliced-in exons, we extracted the splice junction information at both the acceptor and donor sites and classified them as canonical or noncanonical based on the presence of these specific dinucleotide combinations.

To assess reproducibility, filtered transcripts were compared to various independent transcript datasets, including the latest version of GENCODE (v.43), transcripts identified by Workman et al. (*102*), Gao et al. (*103*), Leung et al. (*104*), the Gtex long-read sequencing study (*105*) and transcripts called by CHESS 3 (*106*). This comparison was performed using gffcompare (*107*), and transcripts showing an exact intron chain-match with those from other studies were considered as transcripts observed in those studies.

#### Differential gene and transcript expression, and transcript usage analysis

Differential gene expression (DGE) and differential transcript expression (DTE) analysis between GZ and CP regions were conducted using DESeq2 v1.34.0 (*108*). DGE was conducted using a counts matrix for all expressed genes (n=24,554), controlling for library size through estimated size factors as is standard in DESeq2. Specified covariates included the two regions (GZ and CP) and three biological replicates (donors). The “DESeq” method was then applied with default parameters, and genes with an adjusted p-value cutoff (FDR) of 0.05 were considered differentially expressed.

For DTE, the same procedure was followed, but we only considered transcripts from genes with multiple isoforms and applied a stricter isoform expression threshold, resulting in 102,319 transcripts for testing. In detail, only genes with an average expression of 1 TPM across both regions and isoforms with an isoform fraction [(IF), calculated as isoform expression/gene expression counts] value greater than 0.01 in one or both conditions were retained.

Differential transcript usage (DTU) was performed using DEXSeq (*109*) as implemented within IsoformSwitchAnalyzeR (*110*). The same prefiltered counts matrix as described above for DTE was used, and a donor column was included in the design matrix as a confounding factor. A difference in isoform fraction (dIF) value between the two conditions and an FDR adjusted p-value were calculated for each isoform. We considered isoforms with an adjusted p-value lower than 0.05 as exhibiting DTU, regardless of dIF. At the gene level, the lowest adjusted p-value of any isoform was considered to be the p-value for the gene; therefore genes with at least one significant isoform were considered significant.

#### Alternative Polyadenylation Analysis

The Dynamic analysis of Alternative PolyAdenylation 2 (“DaPars2”) algorithm was used to quantify relative alternative polyadenylation (APA) site usage from bulk fetal tissue IsoSeq data (*45*). Annotated 3’UTRs were extracted from the GRCh37 RefSeq whole gene annotation file from the University of California Santa Cruz (UCSC) Table Browser. SAMtools v1.15 (*111*) was used to calculate read depth of each sample, which was used to normalize sequencing depth difference of samples. DaPars2 was used to align aggregated IsoSeq reads from all samples to 3’UTR regions and calculate the distal polyA usage index (DPUI), i.e. the relative proportion of distal APA site usage compared to proximal APA site usage. DaPars2 first identifies a distal APA site based on where sequencing coverage ends. Identification of a proximal APA site is based on a drop-off point in read density. Finally, DaPars2 adds read counts to quantify usage of the proximal and distal APA sites and calculates DPUI for each transcript. APA events with read coverage less than 10 were excluded. Repeated measures ANOVA accounting for multiple replicates per sample was used to assess how PDUI changed by transcript between GZ and CP samples.

#### Pathway, RNA binding protein target (RBP), and cell-type enrichment analyses

*Pathway enrichment using gProfileR*

The enrichment analysis was performed using gprofileR for Gene Ontology (GO), and Kyoto Encyclopedia of Genes and Genomes databases against all the expressed genes as reference. An adjusted P value cut off of 0.05 was used to determine significantly enriched GO terms or pathways and term size was set to ≤ 1,000 genes. All figures were generated in R using ggplot2.

*RBP target enrichment*

A list of known human RBPs compiled from the PFAM database and interactome capture was downloaded from Sundararaman et al. Table S1 (*112*) and plotted against geneExpr, isoExpr and isoUsage network dendrograms.

RBP target gene sets were gathered from published datasets of brain-enriched RBPs, or downloaded via the web portal (www.encodeproject.org) for ENCODE eCLIP RBPs. See **Table S7** for citations and metadata.

For the set of brain-enriched RBP targets, we downloaded supplemental files of genes identified as containing splicing (AS) or gene expression (GEX) changes after knockdown or knockout and those found through direct binding (CLIP). We chose relevant brain-enriched RBP target datasets using the following criteria – a) the RBP is expressed predominantly in neurons, or has differential expression between neural progenitors and neurons, b) knockdown/knockout or binding of the RBP was assayed in its relevant cell type (e.g. *Rbfox1* knocked out in neurons), c) the RBP is expressed in our isoform sequencing data. For the ENCODE eCLIP dataset, we chose all RBPs that a) are expressed in our isoform sequencing data, b) were assayed in the HepG2 liver cell line and c) have at least two replicate samples.

When applicable, we filtered target lists using quality-control or p-value criteria provided in the relevant publication. For datasets generated from mouse tissue, we kept genes that had one-to-one human-to-mouse homology. For the set of ENCODE RBP targets, we downloaded unfiltered eCLIP bed files for RBPs described in Van Nostrand et al. from the ENCODE portal and applied filtering criteria as described in the paper (peaks with *P*<0.001 and log2 fold-change ≥3 kept).

Direct binding datasets (CLIP for brain-enriched or eCLIP for ENCODE) that were provided as genomic coordinates were overlapped with known gene coordinates using the GRanges R package and the relevant genomic annotation. For datasets generated in mice, we used *mm9* or *mm10* depending on the dataset, and for human datasets we used *hg38*. We required that the coordinates for each (e)CLIP peak had at least 50% overlap with gene coordinates to assign it to that gene.

After QC and filtering steps, we performed enrichment analyses of the overlap between each RBP target set and either a) DGE, DTE and DTU gene lists from bulk isoform sequencing or b) module-associated genes or isoforms from each of the geneExpr, isoExpr and isoUsage networks. Enrichment was performed using a one-sided Fisher’s exact test followed by FDR-correction for multiple testing (adjusted *P*<0.05). We used genes detected in bulk isoform sequencing as the background gene set, except for overlaps using datasets generated in mice, where we used one-to-one homologs detected in bulk isoform sequencing.

*Cell type enrichment*

We used cell type marker genes of the mid-gestation neocortex as previously defined(*3*) (Table S4, “cluster enriched genes”). Briefly, these genes were identified as differentially expressed in one cluster as compared to all other cell clusters in the dataset, and are expressed in at least 10% of the cells of the cluster. Enrichment of cell type markers with a) DGE, DTE and DTU gene lists from bulk isoform sequencing or b) module-associated genes or isoforms from each of the geneExpr, isoExpr and isoUsage networks were performed using Fisher’s exact test followed by FDR-correction (adjusted *P*<0.05). We used all genes detected in (*3*) as the background gene set.

#### Proteomic Analyses

For ORF prediction, we used TransDecoder v5.5.0 (https://github.com/TransDecoder/TransDecoder/) to predict the open reading frame (ORF) for each novel transcript. Local alignment using blastp (*113*) was performed against the reference proteome to identify homologs in UniProt. PFAM domains for all possible ORFs were predicted using the hmmscan tool from hmmer v.3.3.2 using default parameters (*114*) . The best predicted ORF(s) for each novel transcript were selected by TransDecoder on the basis of coding potential, significant sequence homology (blastp) and domain conservation (hmmer) to human proteins. In addition to TransDecoder, CPAT(*115*) was also used to further confirm the coding potential of each novel transcript. With all predicted ORFs, we further performed nonsense-mediated mRNA decay (NMD) analysis by searching for a premature termination codon located more than 55 nucleotides upstream of the last exon-exon junction.

For peptide search and novel protein validation, raw proteomics datasets were obtained from BrainSpan (ProteomeXchange: PXD005445) (*44*) and PsychENCODE (Synapse: syn26017684), and then converted to mzXML format. Spectra search was performed using Comet (release 2019.01 rev. 5) against a protein sequence database with 104,706 human protein sequences from Gencode supplemented with 152,296 coding sequences derived from novel transcripts. Given the sample preparation protocols and the high-resolution mass spectrometry instrumentation settings, we used the following parameters for database search, as per best standards in the field. The precursor ion mass tolerance was set to 20 ppm to account for the error range of the mass analyzer, and the fragment ion bin width was set to 0.02. Semi-digestion by trypsin was used, with up to 3 missed cleavages allowed to maximize the spectra search space within computationally-feasible timelines. Methionine oxidation was set to variable modification. Cysteine carbamidomethylation was set to be static modification. For PsychENCODE proteomics data, peptide N-terminal and lysine (K) tandem mass tag (TMT11plex) were also set as static modifications. After the spectra search, peptide-spectrum-matches (PSMs) were pooled and re-scored using Percolator (v3.05.0) (*116*). PSMs with re-scored FDR-corrected P values < 0.05 were kept. If a peptide from a PSM uniquely mapped to novel ORF(s) but not to GENCODE known protein sequences, the corresponding novel ORF(s) were classified as “peptide-supported”.

#### Weighted gene correlation network analyses (WGCNA)

Network analysis was conducted with the R package WGCNA (*61*) using bulk GZ/CP gene expression, transcript expression, and isoform fraction quantifications. Gene and isoform (transcript) expression were log_2_TPM normalized. Scale free topology was best achieved when adjacency matrices were raised to the soft-threshold powers of 14, 9, and 14 for gene expression, isoform expression, and isoform fractions, respectively. Signed networks were then generated, and modules were identified using the cutreeHybrid function with the following parameters: deepSplit=4, minClusterSize=100, pamStage=FALSE, and cutHeight = 0.999999. To define the biological processes associated with modules, we calculated the module eigengene (first principal component) and evaluated its correlation to known attributes of the dataset including cortical region (GZ/CP). We also conducted over-representation analysis to look at enrichment of each module with known gene lists using a one-sided Fisher’s exact test (cell type and RBP lists) or logistic regression for rare-variant associated disease gene lists, controlling for gene length and coding length. In addition, gene ontology analyses was performed on all modules as described above. Module hub genes and their connectivity were visualized using the igraph package in R.

#### Single cell Iso-Seq (scIso-Seq) library preparation, sequencing, read mapping and QC

Single-cell PacBio IsoSeq libraries were prepared from previously captured (*3*) single-cell full length cDNA generated following the Drop-Seq protocol v.3.1. Cell and UMI barcoded cDNAs from approximately 5,000 single cells were selected from a pool of 40,000 cells and subjected to a second round of PCR amplification for 6 additional cycles with SMRT primer using Kapa Hi-Fi polymerase to generate sufficient cDNA for PacBio library preparation. The amplified product was purified by AMPure beads following the manufacturers’ guidelines. The quality and concentration of the final purified product was assessed using a Bioanalyzer 2000 and Qubit. This purified product was used directly for PacBio SMRT bell template preparation and sequencing. Sequencing was performed on both Sequel I (6 SMRT cells at the UC Davis Genomics Core) and Sequel II machines (7 SMRT cells at the University of Maryland Genomics Core).

Raw IsoSeq subreads were processed using the ccs (CCS v4.0.0) function of PacBio Smartlink v8.0.0 package to generate circular consensus sequences (ccs) from subreads with the following parameters, in addition to the default ones (parameters: --minLength=50 -- minPasses=1 –min-rq=0.8 --min-snr=2.5 ). To ensure selection of full-length isoforms, CCS reads were then scanned and filtered for the presence of 3’ and 5’ adapters using lima function (v1.10.0) running in “isoseq” specific mode. Thereafter, full-length non-chimeric (FLNC) reads were extracted using the isoseq3 refine package (v3.2.2; parameters: --min-polya- length 20 --require-polya). Following this extraction step, Across all 13 SMRT cells, over 19.1 million full-length non-chimeric reads were retained. To control for PCR amplification biases in single-cell data, FLNC read sequences were then collapsed by UMI and cell barcodes. The fastq sequences were mapped to GRCh37 human genome reference using minimap2 (version 2.17-r954) with parameters “--ax splice -uf --secondary=no -C5 -O6,24 -B4 --MD -t 32”. We then used TransciptClean using parameters “--maxLenIndel=5 --maxSJOffset=5 --primaryOnly” to correct for reference-based errors in long-read alignments including mismatches, indels, and non-canonical splice-sites.

#### scIso-Seq transcript discovery and quantification

To select high-quality cells for all subsequent analyses, we used a cumulative distribution function (knee plot) and retained cells with >85 UMI reads per cell as indicated by the inflection point in the plot (**Fig. S9**). To leverage the high-depth sequencing and clustering resolution of a 40K cell dataset, we matched long-read sequenced cells with those from short-read sequencing ((*3*)) using their unique cell barcodes and obtained all associated metadata. An independent sam file was generated for each of the matched cells with the read output from TranscriptClean and a config file was generated for input into TALON. TALON module was run on the config file along with the TALON database created for the bulk GZ/CP Iso-Seq data with parameters “--cov 0.85 --identity 0.8”.

Similar to bulk Iso-seq, a GTF annotation file was generated for isoforms passing the following criteria:

(1) isoform labeled as known;

(2) isoform labeled as ISM/NIC/NNC/Genomic/Antisense/Intergenic ***and*** supported by 3 independent reads, ***and***

(a) present in two independent donors ***or***

(b) have independent support for 5’ end within 100bp upstream or downstream and for 3’ end within 200 bp upstream or downstream of the start and end position of the transcripts ***or***

(c) was reported in an independent long-read isoform database

For support analysis of the 5’ end, we used CAGE peak annotations from the FANTOM5 database (*40*) and assessed overlap using bedtools. The genomic position of the 5’ end for each isoform was obtained from our GTF file and then checked for overlapping CAGE peak within 100 bp region immediately up or downstream of the 5’ end position by using bedtools. For the 3’ end support, we used the published polyA-site database (*41*) as well as computationally-predicted polyA motifs as implemented in TALON (*34*) . The resulting whitelists were used to generate filtered abundance files for transcript quantification as well as custom-filtered GTF annotations for subsequent studies. We used bedtools tools to identify spliced-in exons not present in the reference transcriptome (**Table S5B**).

#### scIso-Seq clustering, differential expression, and lineage trajectory inference

Single-cell isoform count matrices, generated by the TALON abundance module, were imported into Seurat (v3). To avoid potential cell multiplets or low-quality cells, we removed cells with over 2,000 unique isoform counts, and those for which the total mitochondrial reads exceeded 7%. We then filtered out isoforms not expressed in a minimum of 3 cells. To correct for technical variance associated with single-cell methods, including differences across cells in read-depth and gene detection, we performed negative binomial variance-stabilizing transformation (VST) on the filtered count dataset using sctransform (*117*). To correct for potential differences in cell health, we regressed out mitochondrial gene expression using vars.to.regress in sctransform. Finally, to maximize biologically-relevant information, we set “variable.features.n” to 10,000 which uses the top n variable genes for clustering. To minimize the effect of technical noise on clustering, dimensionality reduction was performed by principal component (PC) analysis. The first 26 PCs were selected based on elbow plot analysis as implemented in Seurat. UMAP embeddings were calculated with “RunUMAP” function using the first 26 PCs, 15 neighboring points, and a minimum distance of 0.15. To examine the impact of gene and isoform expression on single-cell clustering, we first used the cell identity from a previous short-read sequencing-based analysis ((*3*) ) with UMAP cell embeddings from the present study to generate the gene expression based UMAP plot. For isoform expression based cell clustering, cells were clustered using the “FindNeighbors” and “FindClusters” functions in Seurat with a resolution of 1.6. Thereafter, we used “DimPlot” function to generate the UMAP plot. For the 4,281 cells, clustered by isoform expression with resolution of 1.6, we distinguished major cell types in the UMAP map according to known markers genes associated with the isoforms as well cell type information extracted from (*3*). Stability of identified clusters was evaluated by bootstrapping as implemented in “bluster::bootstrapstability” (*118*). Cells and their isoform-based cell cluster information were provided in Supplementary Table S5A. Differentially-expressed isoforms identified for each cell type were identified by the “FindAllMarkers” function of Seurat (logfc.threshold = 0.25 pos.only = T, min.pct = 0.15) and provided in Supplementary Table S5C. To identify differentially expressed isoforms between specific pairs of clusters, we used the “FindMarkers” function (Supplementary Table S5D). Single-cell trajectory analysis was performed using Monocle3 (v.0.2) (*119*). We first converted the Seurat object into a Monocle3 object and then inputted the Seurat UMAP coordinates and cluster labels into the learn_graph function to generate the trajectory path.

#### scIso-Seq differential transcript usage analysis

To identify transcripts that have different usage across cell types we conducted single-cell DTU analysis (**Table S5E**). We first filtered out the transcripts with fewer than three reads in all cell types and then examined the remaining transcripts individually. For each remaining transcript, we performed an overall differential test and 16 cell-type-specific differential tests (corresponding to the 16 cell types). The overall differential test defines whether the transcript has different usage in at least two cell types, while each cell-type-specific differential test defines whether the transcript has differential usage between the specific cell type and the union of the other cell types. For all tests for a transcript i, we used the binomial regression with the random response variable Yi as the transcript’s read count in a cell, and the fixed parameter “total” ni in the binomial distribution is the transcript’s corresponding gene’s read count in a cell (note that the gene has other transcripts, so Yi<ni). For the overall differential test, we included the following covariates, each with one value in a cell: cell type (the covariate of interest), cell library size (for normalization), and donor (for batch effect removal); for each cell-type-specific differential test, we changed the covariate of interest from cell type to a binary indicator of whether a cell belongs to the specific cell type or not. Then in each test, we tested whether the covariate of interest has a non-zero effect; we ran the binomial regression using the R function glm with family="binomial" in the stats package, and we calculated the p-value (for inclusion vs. exclusion of the covariate of interest) using the Chi-squared test in the R function anova in the stats package. In summary, we obtained one p-value per transcript per test. Finally, for each test, we used the BH procedure to correct all transcripts’ p-values for false discovery rate control.

**RT-PCR validation of novel exons:**

500 nanograms of total RNA was reverse transcribed using SuperScript IV reverse transcriptase (Invitrogen Catalog number: 18090010) and with oligod(T) primer following manufacturer’s recommendation. PCR was used to amplify regions spanning the novel exons and nearby known spliced in/our exons. Primer sets were designed to anneal to the novel exon and to a nearby known exon. Primer sequences can be found in **Table S8**. PCR amplification was performed with Q5 High Fidelity DNA polymerase. PCR products were separated in 1.5% agarose gel.

#### Rare variant enrichment analyses

We conducted a series of enrichment analyses to localize rare-variant association signals from large-scale whole exome and genome sequencing studies of neurodevelopmental and psychiatric disorders within our transcriptome annotation features, GZ/CP differential gene and isoform expression and usage (DGE, DTE, DTU), cell type-specific isoform expression and utilization, and WGCNA networks. Gene lists for each disorder were compiled using the indicated significance threshold appropriate for each study: ASD.fuTADA (*82*) (q-value<0.1), NDD.fuTADA (*82*) (q-value<0.1), BIP.bipex (*85*) (P<0.01, uncorrected), EPI.epi25 (*86*) (P<0.01, uncorrected), SCZ.schema (*84*) (q-value<0.1), DDD.kaplanis (*83*) (Bonferroni corrected P<0.05). ASD.SFARI.1, ASD.SFARI.1or2 and ASD.SFARI.S were generated according to the indicated SFARI gene scoring (<https://gene.sfari.org/database/gene-scoring/>). The manually curated EPI.helbig list was downloaded from: <http://epilepsygenetics.net/wp-content/uploads/2023/01/Channelopathist_genes_internal_2023_v2.xlsx>

For transcriptome annotation features, enrichment was calculated using logistic regression controlling for gene length and coding length [number of transcripts per gene (n_transcripts_log2), number of exons per gene (n_exons_log2) and genes containing novel exons (novel_exon)], or controlling for gene length, coding length, and transcript expression (n_transcripts_log2|tpm). Logistic regression controlling for gene length and coding length was used to assess enrichment for DGE/DTE/DTU (Table S3), co-variation modules, and cell type isoform expression (Table S5C). Background was set to all protein-coding genes. Significance threshold was set to FDR-corrected pvalue<0.05 across all tested features.

#### Re- annotation of variant consequences

*De novo* variants from ASD (*87*–*89*) and intellectual disability/developmental disorders (*89*) were collected from the literature , and the variant effect was annotated by running Ensembl VEP (v108.2) with the --most_severe flag. We performed two rounds of annotation, the first one using GENCODE v.33 GTF file and the second one by supplementing the previous annotation with predicted ORFs from newly identified transcripts.

#### Prediction of cryptic splice variants

De novo variants from ASD (*87*–*89*) and intellectual disability/developmental disorders (*89*) were collected from the literature, and their effect on splicing was predicted by SpliceAI (*23*). A gene annotation file containing both Gencode (v33) transcripts and novel transcripts from this study was used as input for SpliceAI, while all other settings were kept as default. △Score > 0.1 was used to define potential cryptic splice mutations. Cryptic splice mutations may have effect on more than one transcript. Cryptic splice mutations which have effect on at least one Gencode transcript were considered to be annotated based on Gencode. Cryptic splice mutations which have effect exclusively on novel transcripts were considered to be annotated based on the expanded annotation containing our novel transcripts.

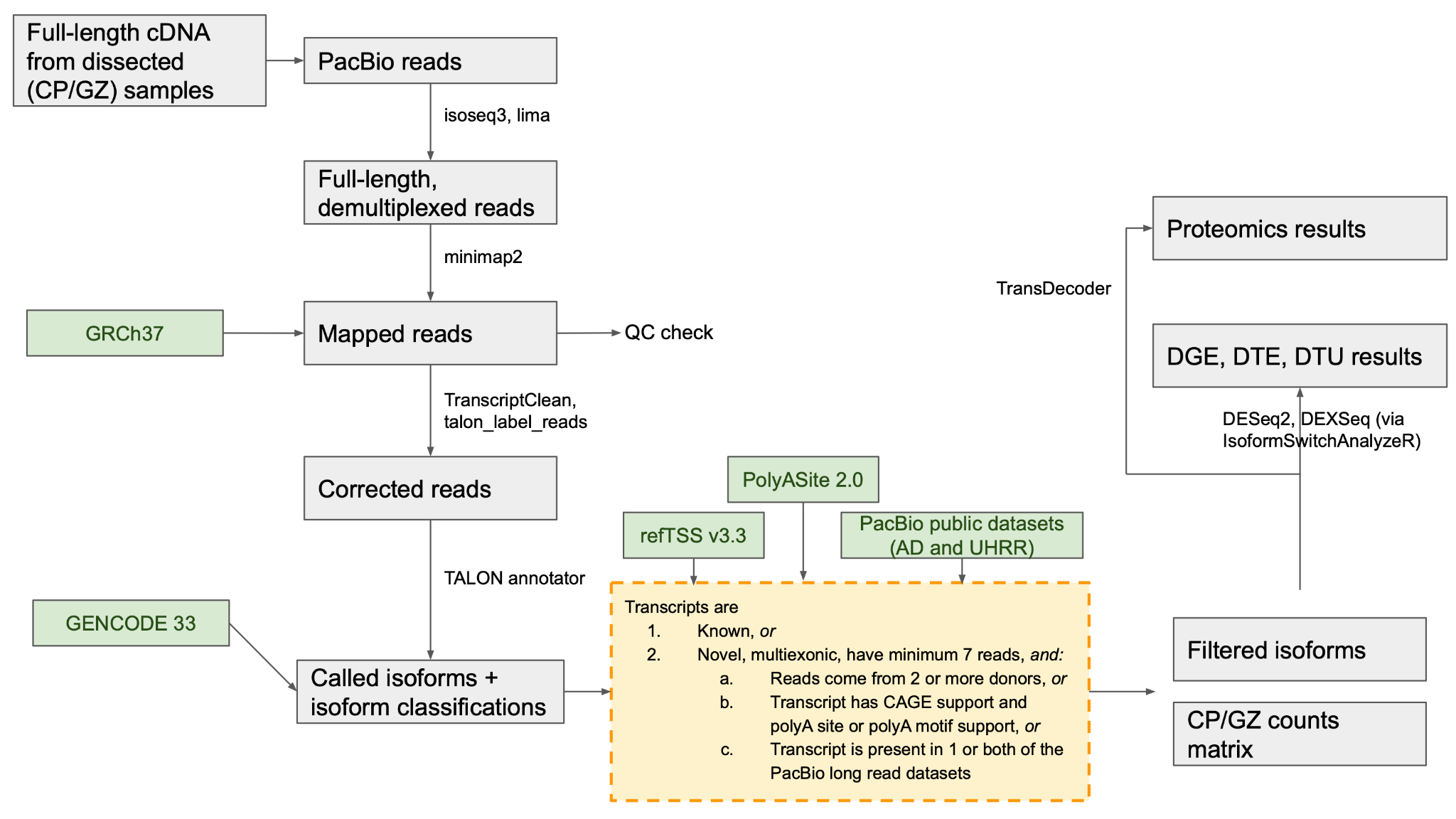

#### Fig. S1. Detailed experimental workflow, data generation, and quality control for bulk tissue IsoSeq.

Briefly, full length isoform sequencing data was generated from CP and GZ region of human fetal brain from three donors. Quality filtered reads were mapped to the human reference genome GRCh37 with minimap2 followed by error correction in the aligned file through TranscriptClean. Isoforms were called on the error corrected aligned data by TALON tools. Called isoforms were further filtered based on independent support at 5’ and 3’ end of the transcripts and minimum support reads (highlighted with the dash-lined square in the figure). Further validation of the filtered isoforms was performed based on proteomic support for transcripts. Differential transcript usage (DTU) across CP and GZ samples was quantified using DEXSeq. Differential transcript expression (DTE) and differential gene expression (DGE) were detected using DESeq2. Downstream visualization and categorization of isoform switching was conducted with IsoformSwitchAnalyzeR.

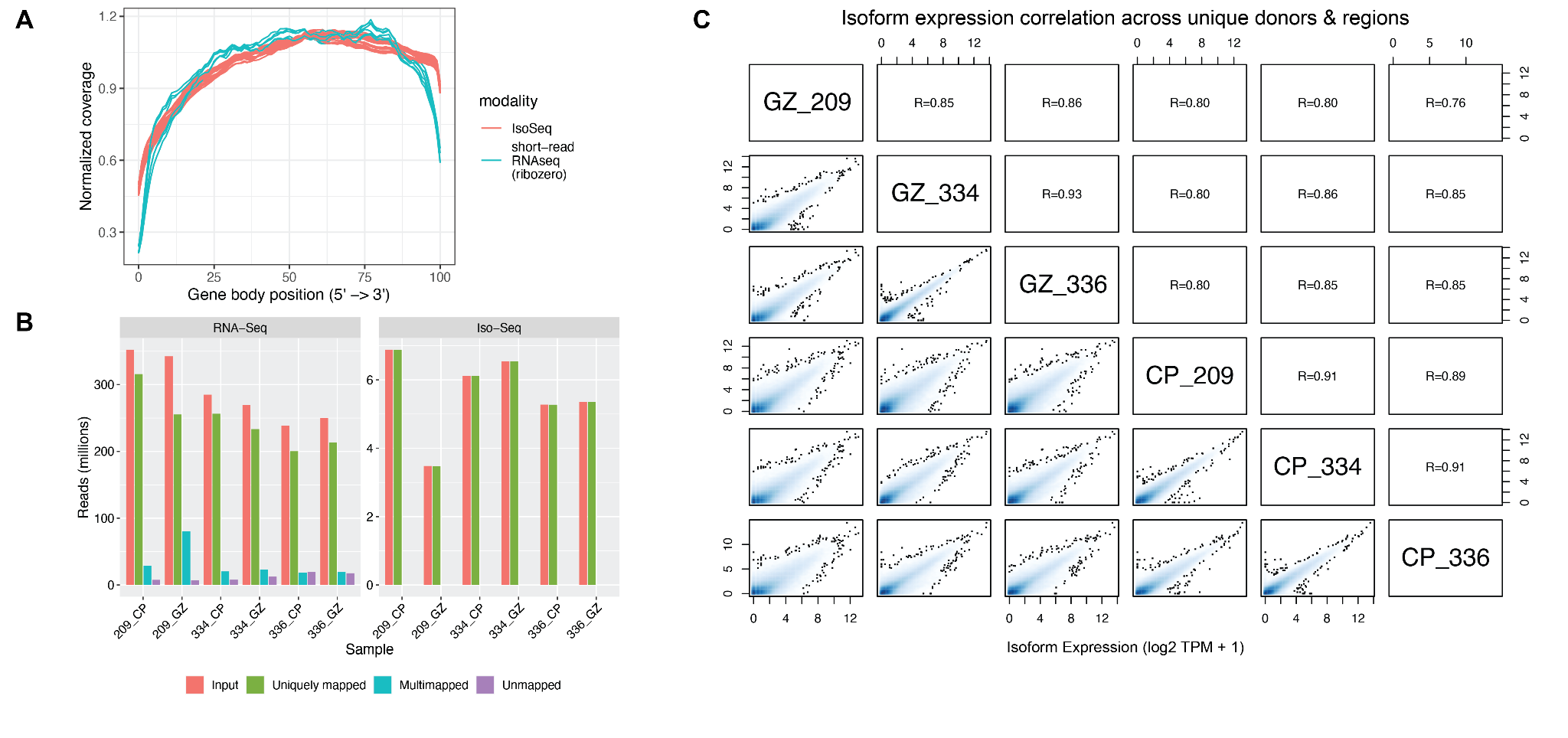

#### Fig. S2. Quality metrics of bulk sequencing data

(**A**) Normalized gene body (5’ to 3’) coverage of matched samples profiled by long-read sequencing (red) and short-read total RNA-Seq with rRNA depletion (blue). (**B**) Number of input reads compared to uniquely mapped reads, multi-mapped reads, and unmapped reads across samples in RNA-Seq and Iso-Seq, after mapping with STAR and minimap2, respectively. (**C**) Isoform-level expression correlations across unique donors and regions. Top panel indicates pearson correlation coefficients.

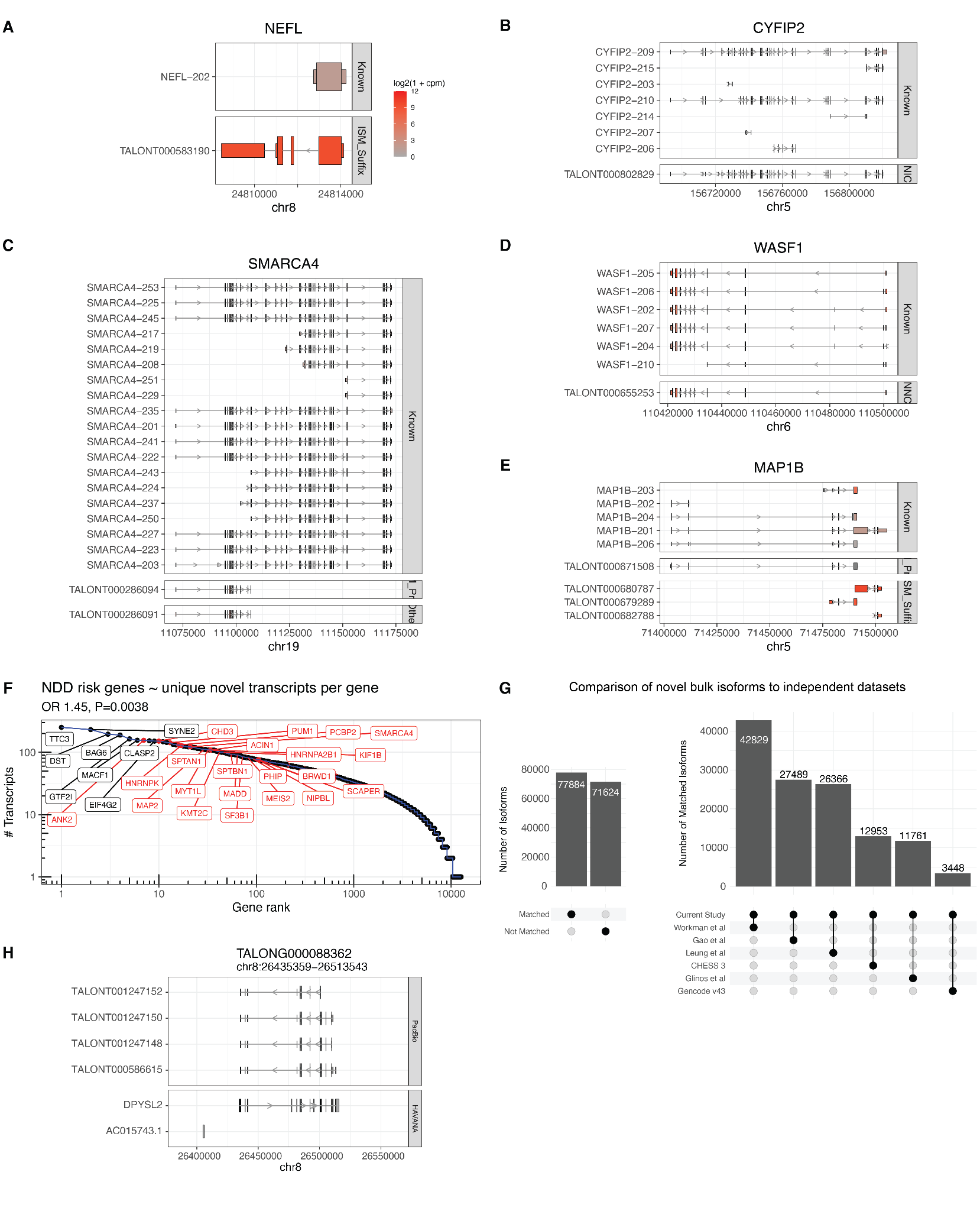

#### Fig. S3. Characteristics of genes with novel transcripts.

(**A-E**) Plots of highly expressed, novel transcripts across categories. Transcripts are colored according to their expression levels as denoted in **fig. S3A**. (**F**) The number of unique, novel transcripts significantly predicts NDD risk genes (red), controlling for total expression, gene length, and coding length. (**G**) Comparison of novel transcripts identified in the bulk transcriptome in this study with those from publicly-available transcript databases and other long read studies. Barplots show the number of novel transcripts detected by bulk Iso-Seq across all studies (left panel) or across specific datasets (right panel). (**H**) A novel multi-exonic, alternative spliced gene is identified on chromosome 8, antisense to *DPYSL2*.

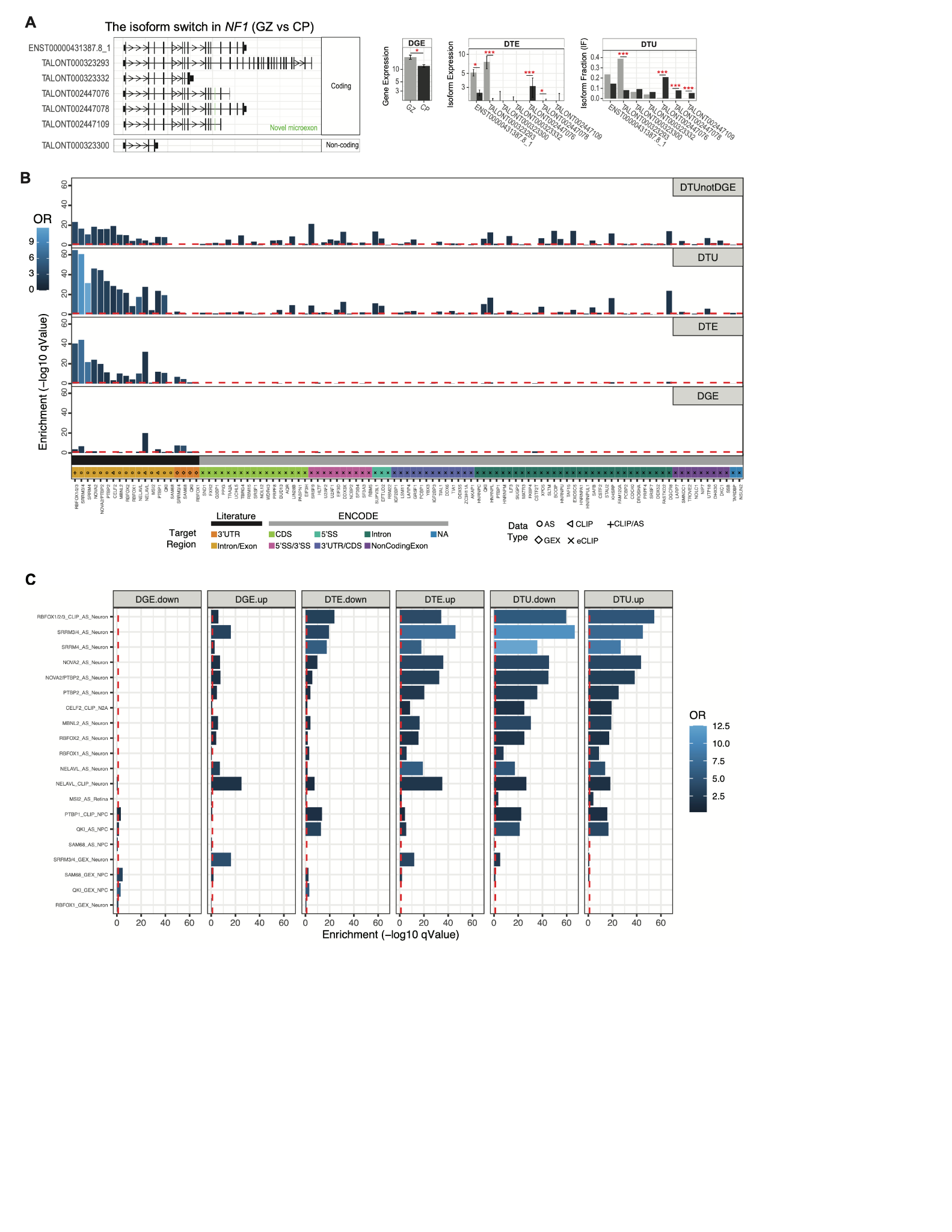

#### Fig. S4. Isoform switching during neurogenesis

(**A**) Switch plots in *NF1.* (**B**) RBP target enrichments for genes containing DTU, DTE or DGE. Odds ratio (OR) is shown in the color bar. RBP target sets are organized by those targeted by known brain-enriched RBPs (yellow, orange bars), followed by RBPs profiled by eCLIP in the ENCODE database and organized by their major target region (e.g. intron- versus exon-binding). (**C**) RBP enrichments for brain-enriched RBPs in genes changing up/down in expression and/or isoform usage. Note gene expression targets of progenitor associated RBPs (PTBP1, SAM68) are more enriched in genes found in the GZ (DGE.down), while targets of neuronal RBPs (NELAVL, SRRM3/4) show enrichment in CP (DGE.up).

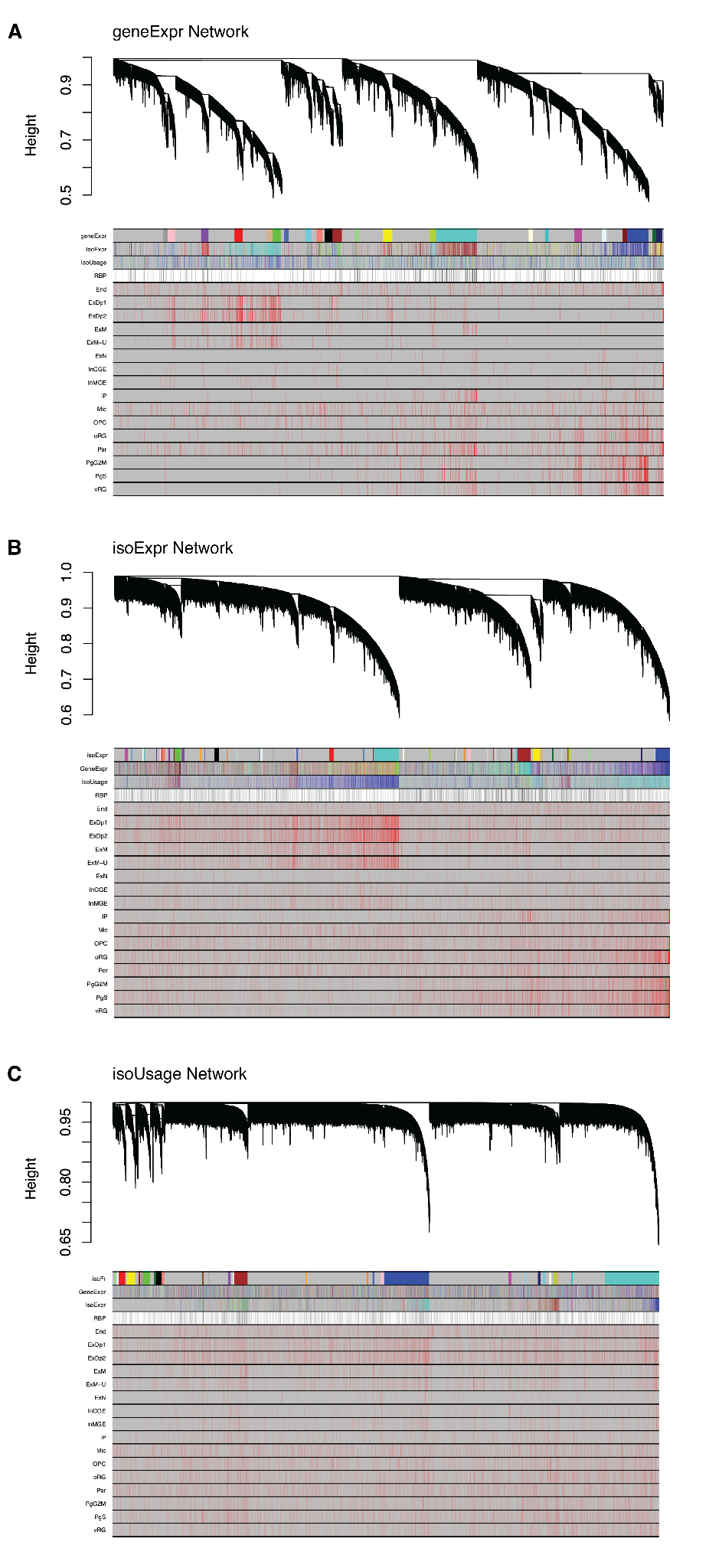

#### Fig. S5. Gene and isoform-level co-expression networks.

Co-expression network dendrograms shown with cell type markers and expression of known RBPs. (**A**) Gene expression network (geneExpr). (**B**) Isoform expression network (isoExpr). (**C**) Isoform usage network (isoUsage).

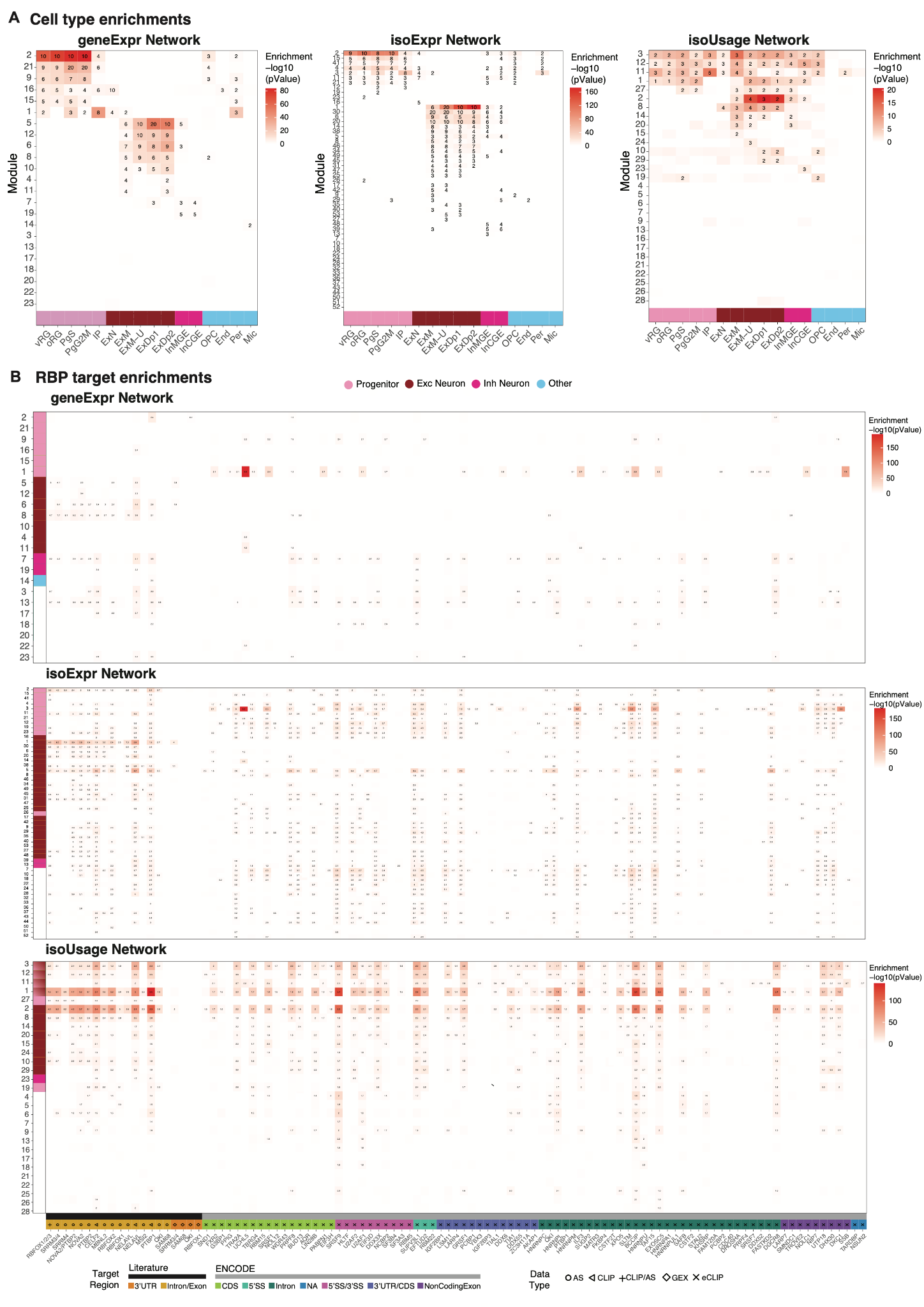

#### Fig. S6. Cell type and RBP target enrichments in co-expression networks.

#### (A) Global cell type enrichments for modules of the three networks. geneExpr and isoExpr modules are strongly driven by cell type, while the isoUsage network is much less so. A set of isoExpr modules is enriched for specific progenitor and neuron subtypes. Among progenitors, isoExpr.M19 is uniquely enriched in PgS, while M23 and M26 are enriched only in oRG. Among neuronal cell types, isoExpr.M16, M9, M27, and M13 are uniquely enriched in ExN, ExM, ExM-U, and InMGE, respectively. Cell classess are indicated by the color bar (bottom), and individual cell types are plotted on the x-axis.(B) Global RBP target enrichments for the same network modules as above. Modules are organized on the y-axis by cell type as in (A), and RBP target sets are organized by those regulated by brain-enriched RBPs (black) or those profiled in the ENCODE dataset (grey). Additionally, ENCODE RBP targets are organized by their target region.

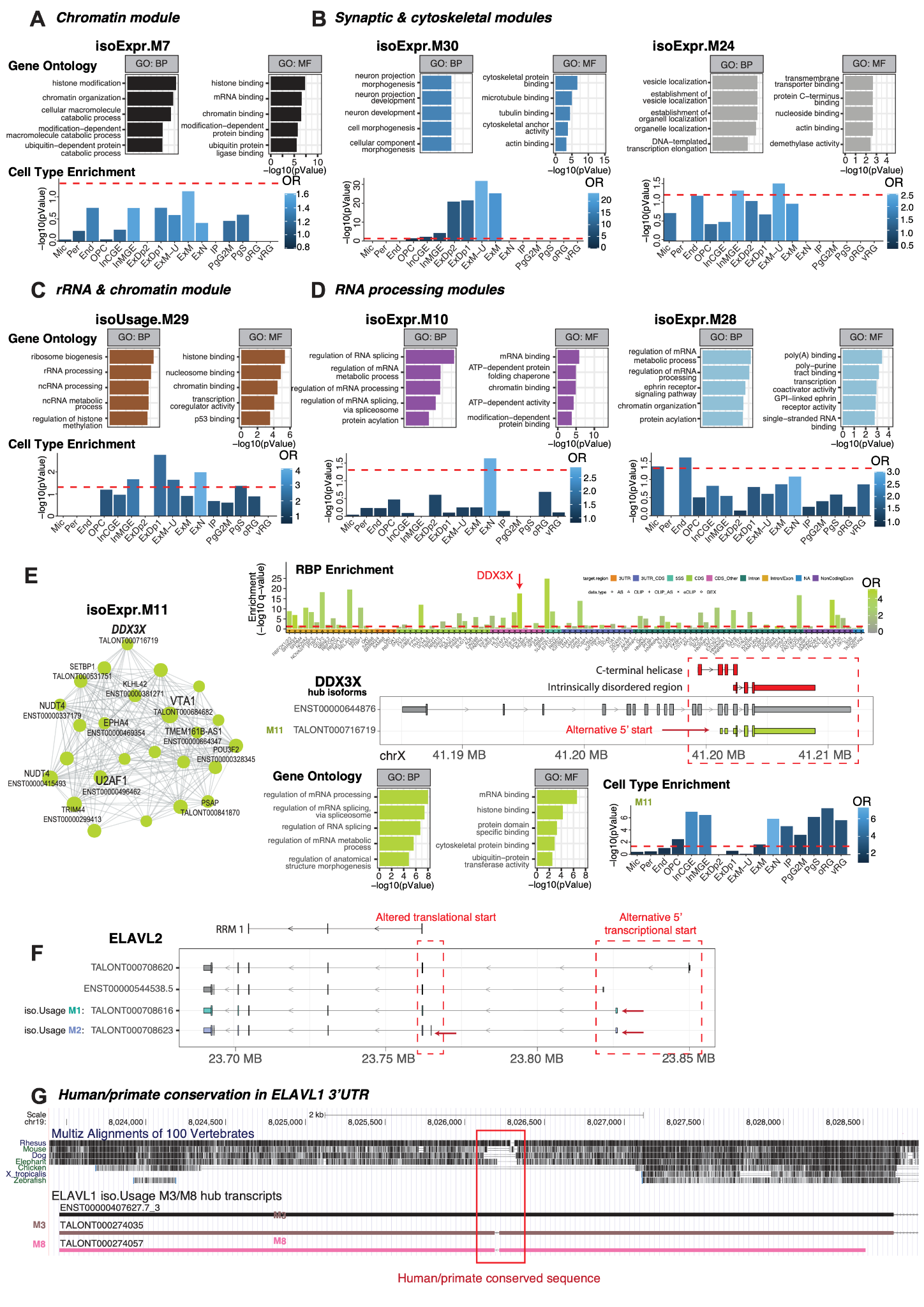

#### Fig. S7. Shared gene ontology processes and disease enrichment in co-expression network modules.

Highlighted isoExpr and isoUsage modules with shared gene ontological processes and significant NDD, DDD and/or ASD gene enrichment. (**A**) isoExpr.M7 is a chromatin regulation module broadly used across cell types and enriched in NDD, DDD and both non and syndromic ASD. NDD and ASD genes are enriched in (**B**) excitatory neuron synapse modules isoExpr.M30 and M24 and (**C**) neuronal ribosomal RNA and chromatin module isoUsage.M29. (**D**) Along with isoExpr.M11, isoExpr.M10 and M28 are enriched in NDD, DDD and non-syndromic ASD and regulate multiple aspects of RNA processing. (**E**) The isoExpr.M11 module is driven by an isoform of the ASD gene *DDX3X* and enriched in RNA processing and cytoskeletal genes. Left, module plot highlighting hub isoforms. Top, RBP analysis reveals strong *DDX3X* target enrichments. Middle, transcript models for the *DDX3X* hub isoform. Box and arrow highlight an alternative 5’ start in the hub transcript resulting in a protein predicted to have no helicase activity but retaining an intrinsically disordered region. Bottom, biological and molecular function GO terms, and cell type enrichments encompassing progenitors and early neurons. (**F**) Transcript models of the *ELAVL2* hub transcripts in iso.Usage M1 and M2. The M2 isoform includes exon 2, which adds 29 amino acids to the first RNA-binding motif (RRM 1). Non-hub transcripts (gray) are presented as examples of alternative 5’ transcriptional starts. (**G**) UCSC Genome Browser view the *ELAVL1* 3’UTR. A ~200bp sequence conserved across human and primates (red box), is not used in iso.Usage M3/M8 *ELAVL1* hub transcripts.

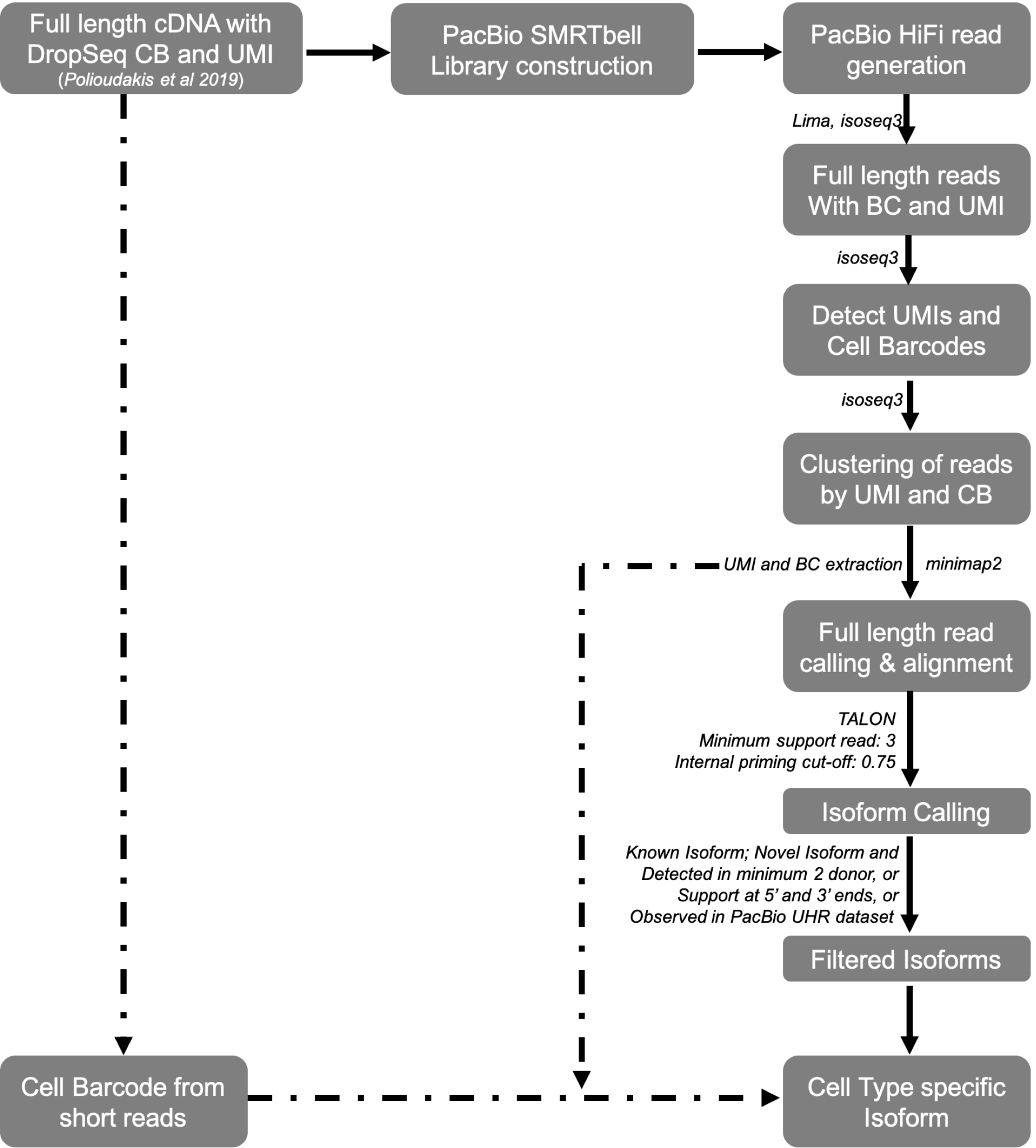

#### Fig. S8. Detailed experimental workflow, data generation, and QC for single-cell isoform sequencing.

Amplified, full-length (pre-tagmentation) cDNA libraries with incorporated Cell Barcodes (CBs) and Unique Molecular Identifiers (UMIs) were obtained from ~5000 single cells isolated from GZ and CP regions across 3 unique donors aged PCW15-16 and were used for SMRTbell library construction and sequencing. Full-length, high quality (HiFi) circular consensus sequencing (CCS) reads were retained only if they contained a polyA sequence of at least 20bp, as well as 3’ and 5’ adapters. CB and UMI sequences were extracted from the filtered full-length reads via isoseq3 tag module and collapsed by UMI and cell barcodes to generate de-duplicated reads. CBs detected by isoseq3 were matched with CBs from high-depth short-read scRNA-Seq previously used to define cell clusters. Reads were mapped to the human reference genome (GRCh37) using minimap2. Putative alignment errors were corrected by TranscriptClean. Thereafter, TALON was used to catalog isoforms from the aligned reads using Gencode v33 as the reference transcriptome. Putative novel isoforms called were further filtered based on several criteria including: multi-exonic structure, identification across multiple independent donors, support from 3+ deduplicated full-length reads, as well as external 5’ and 3’ end support and/or identification in external reference databases.

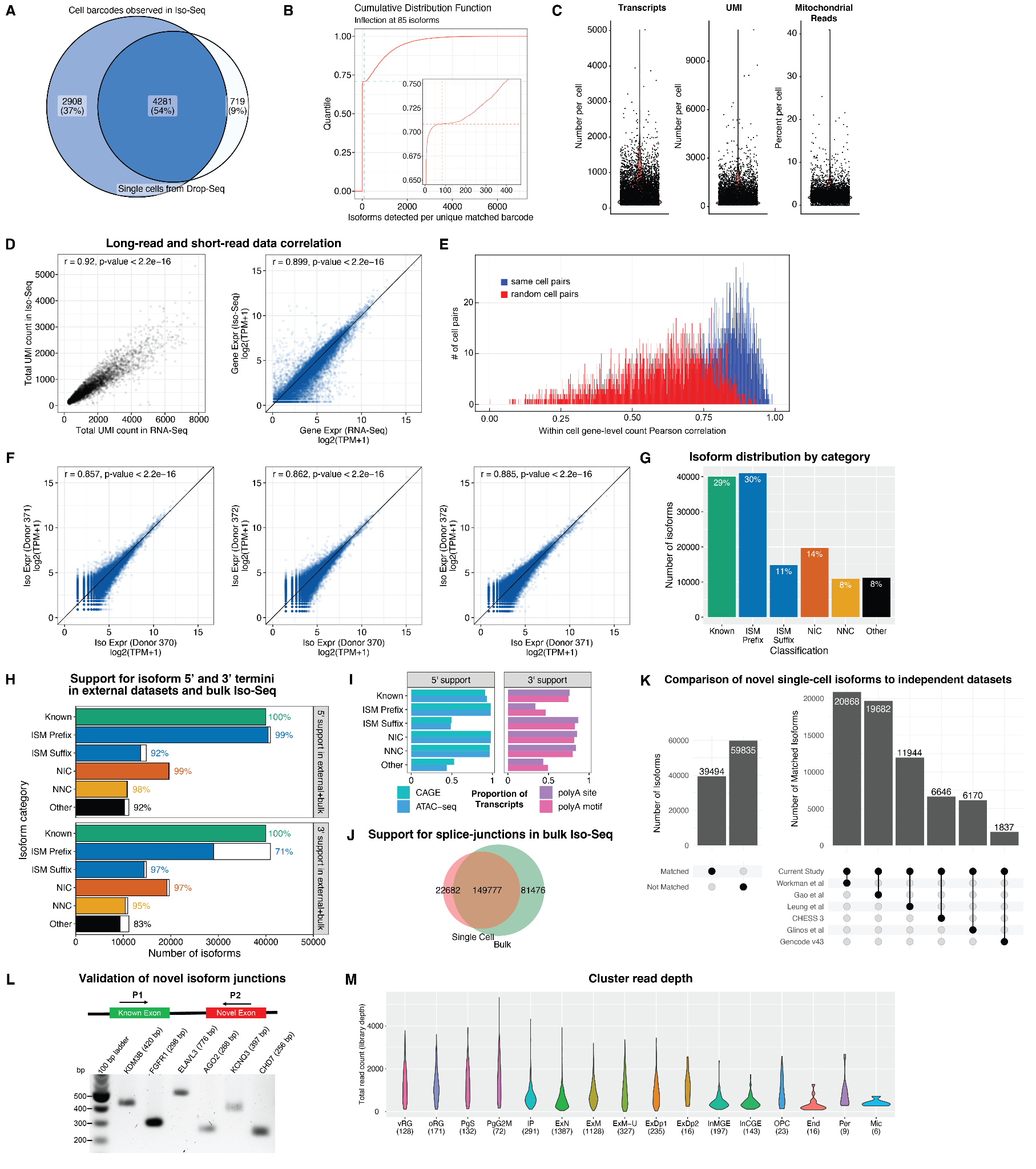

#### Fig. S9. Processing, QC, and validation of single-cell long read sequence data (scIso-Seq).

(**A**) Venn diagram showing cell barcode overlap between cells sequenced by short-read and long-read sequencing following capture by Drop-seq. Approximately 5,000 cells from a pool of 40,000 cells captured by Drop-seq (*3*) were sequenced by long read sequencing, of which 4,281 (86%) were detected by scIso-Seq. An additional 2,908 cells, although passing QC filters, do not match cell barcodes in short read sequencing and were not used in subsequent studies. (**B**) Cumulative distribution function (knee plot) demonstrating the cell barcode count cutoff for identification of true single cells over background. Inset shows magnified section at the inflection point. (**C**) Violin plots displaying the number of isoforms (530, 449), the number of UMIs(757, 607), and percentage of mitochondrial reads(2.2, 2.06) per cell, demonstrating isoform detection in line with single-cell approaches and low mitochondrial reads consistent with high cell health. Mean and median, respectively indicated within parenthesis. (**D**) Isoform detection (left) and gene expression (right) correlation between scIso-Seq (long read) and single-cell 3’ end sequencing (short read) (*3*). Each data point represents a cell. Detection and expression level of transcripts are both highly correlated between sequencing methods. (**E**) Pearson correlation of gene expression between short-read and long-read sequencing, demonstrating high correlation for matched cells (blue) but not for random pairs of cells (red). (**F**) Transcript expression correlation across all donors profiled, demonstrating high inter-donor reproducibility. Each dot represents the transcript expression across each pair of donors. (**G**) Distribution of total isoforms detected by catalog annotation categories. (**H**) Independent validation of detected scIso-Seq transcripts with bulk tissue (CP/GZ) Iso-Seq-detected transcripts and transcripts from 6 independent datasets , demonstrating broad support for most isoform termini. (**I**) External validation of isoforms by independent datasets as explained in **Fig 2B**. (**J**) Venn diagram showing the overlap between splice junctions detected in scIso-Seq and bulk tissue Iso-Seq. ~87% of scIso-Seq-observed junctions are present in bulk Iso-Seq. (**K**) Comparison of novel transcripts identified by scIso-Seq of developing human neocortex with publicly-available transcript databases and other long-read studies. Barplots show the number of novel transcripts detected by scIso-Seq across all studies (left panel) or across specific datasets (right panel). (**L**) Validation of novel exon junctions by RT-PCR from mid-gestation cortex. Expected size is shown in parenthesis along the name of the gene. Regions were amplified as described in Fig 2E and Methods. (**M**) Violin plots sh­owing a balanced number of filtered reads across all clusters. Number of cells per cluster within parenthesis.

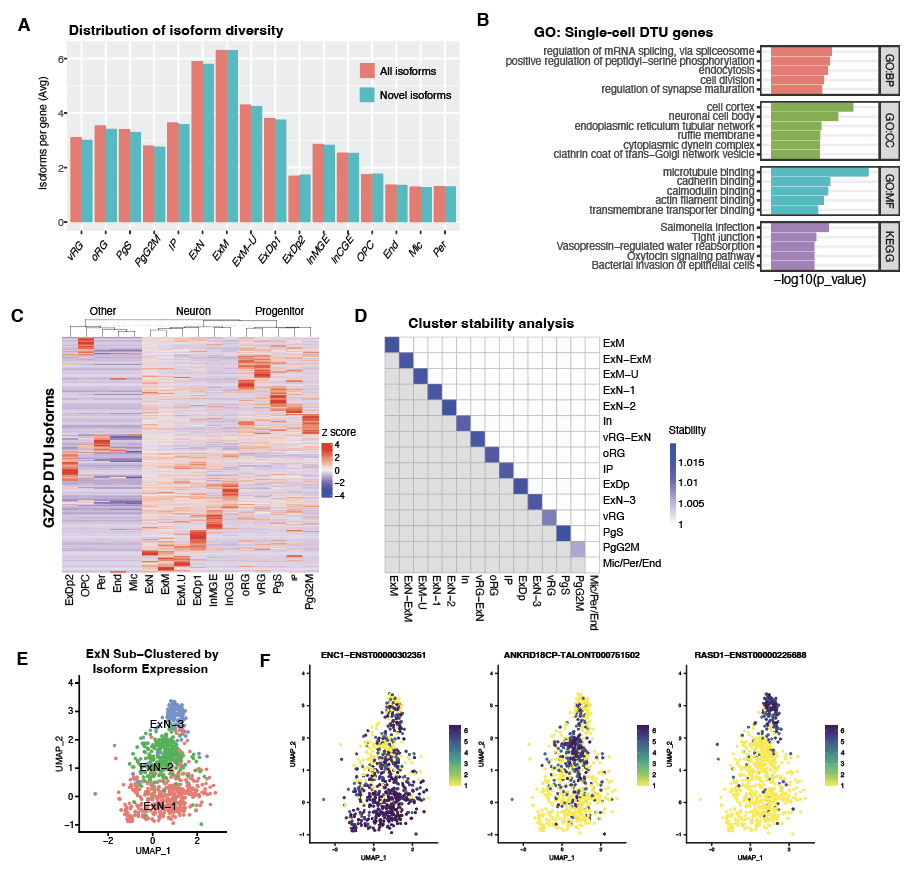

#### Fig. S10. Isoform diversity and utilization, and isoform-based cell clustering.

(**A**) Distribution of isoforms per gene across cell types in the developing human cortex shows ExN and ExM neurons exhibit the highest isoform diversity. “All isoforms” are normalized by all expressed genes in the respective cell type, whereas novel isoforms are normalized to novel genes expressed. (**B**) Gene ontology analysis of cell type-specific DTU genes (**C**) Heatmap of GZ/CP DTU isoform expression across individual cells. GZ/CP DTU isoforms cluster by major cell classes. (**D**) Cluster stability analysis for isoform-based clustering demonstrates high cluster reproducibility. (**E**) and (**F**) Isoform-based sub-clustering of ExN subtypes along with dominant isoform expressed in each sub-cluster.

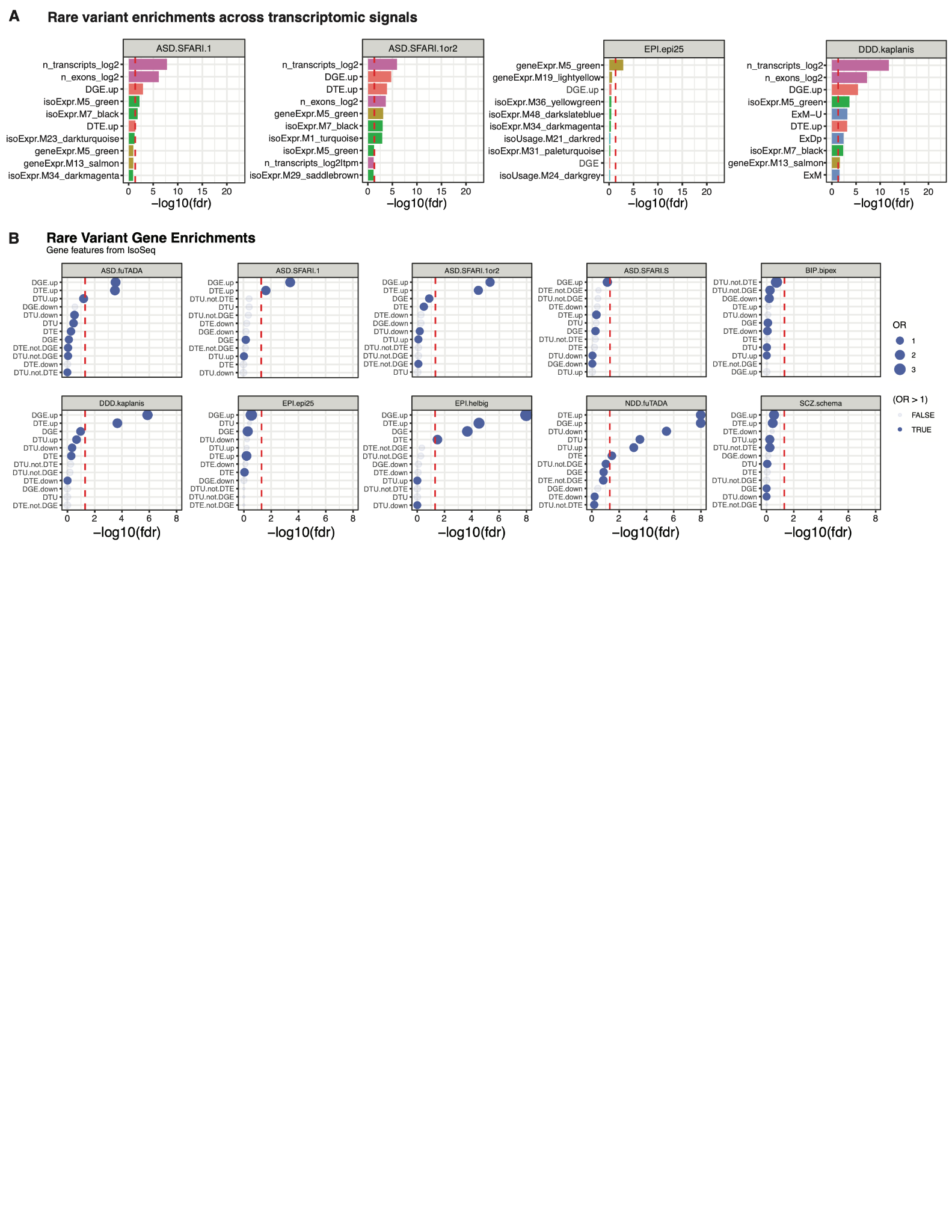

#### Fig. S11. Enrichment across transcriptomic signals of rare variant risk genes for neuropsychiatric disorders.

**(A**) Enrichment of transcriptomic features, differential expression analyses across cortical regions and cell types or isoform expression and usage networks with neuropsychiatric disorders. Red line indicates the FDR-corrected (across all tested features) significance threshold. (**B**) Enrichments for differential gene and isoform expression or utilization across GZ/CP cortical regions. Red line indicates the FDR-corrected (across shown comparisons) significance threshold.

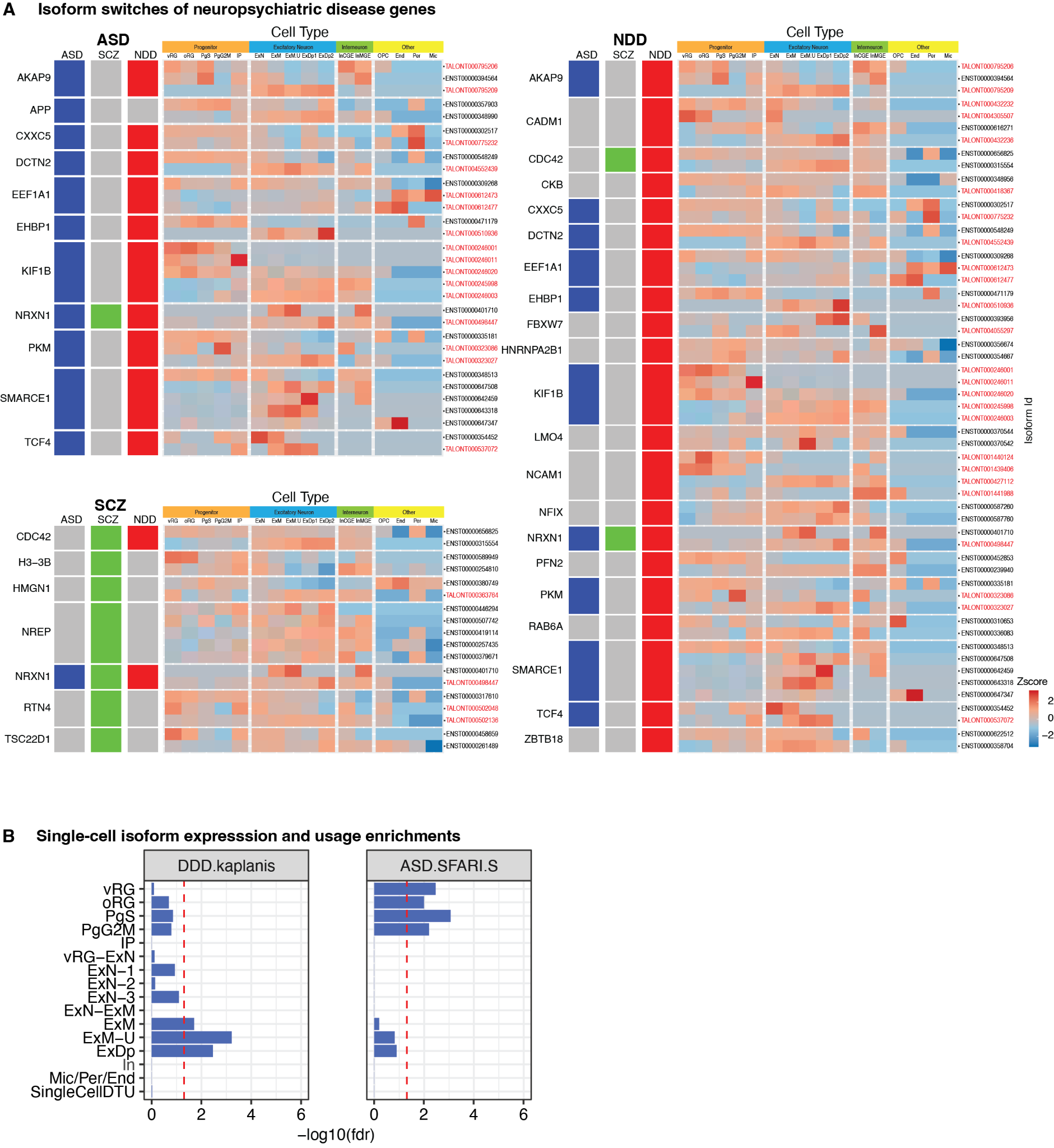

#### Fig. S12. Differential isoform usage of disease risk genes across cells in the developing neocortex.

(**A**) Heatmap of isoforms from NDD (*83*), ASD (*82*) and SCZ (*84*) risk genes showing differential usage across the cell types of the developing cortex. Isoforms from risk genes with two or more DTU isoforms (p.adj <0.05) were plotted. Color bar indicates the average Z score across the entire cluster. Novel isoforms are labeled in red. (**B**) Cell type enrichments indicate differential isoform expression and utilization in NDD and ASD.

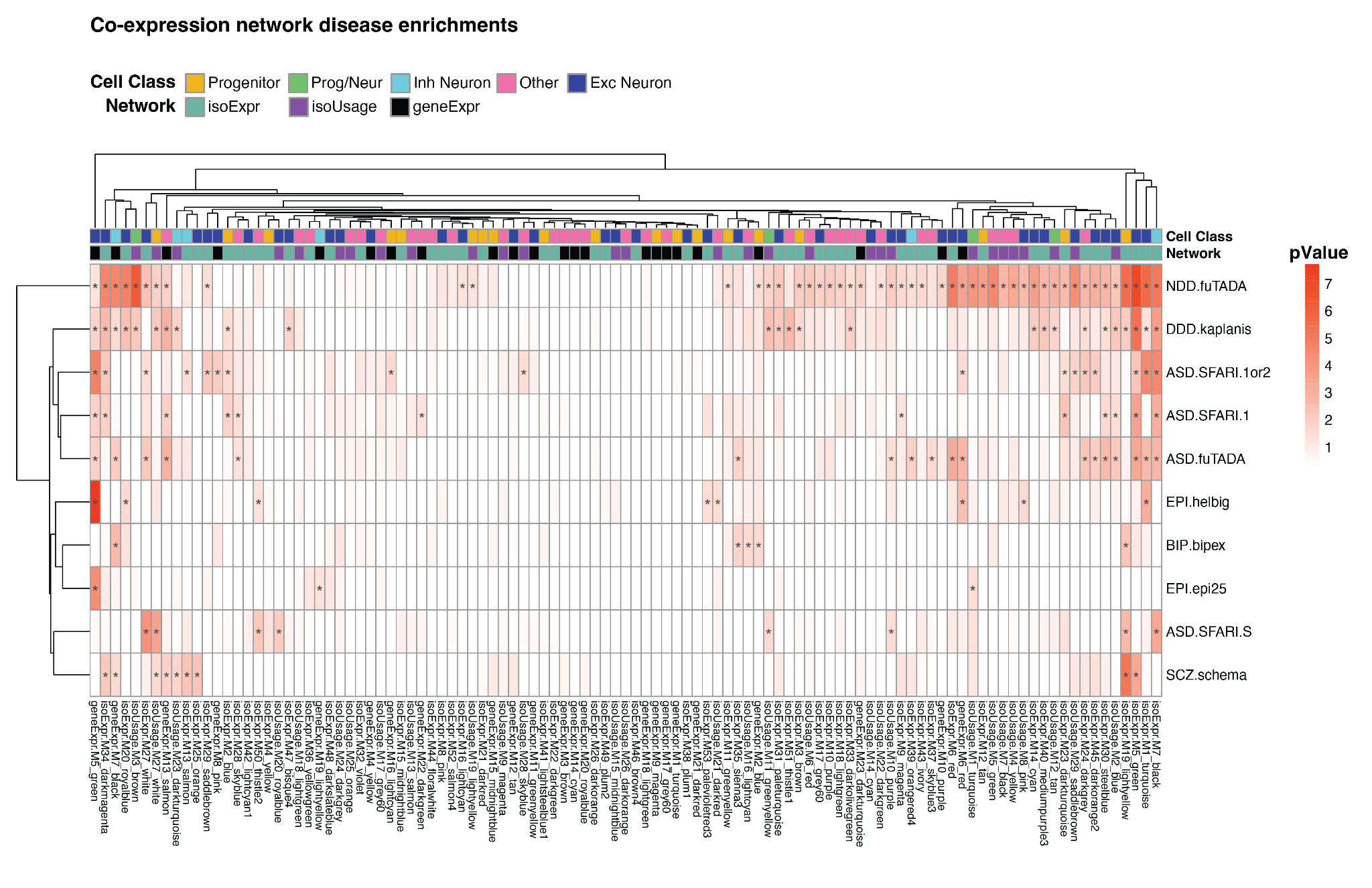

#### Fig. S13. Co-expression networks across neuropsychiatric disorders.

Enrichments of geneExpr, isoExpr and isoUsage networks across neuropsychiatric disorders. Of 105 disease gene-enriched modules (pnominal<0.05), 51.9% were isoExpr (53/105), followed by 26.8% isoUsage (29/105) and 21.3% geneExpr (23/105).

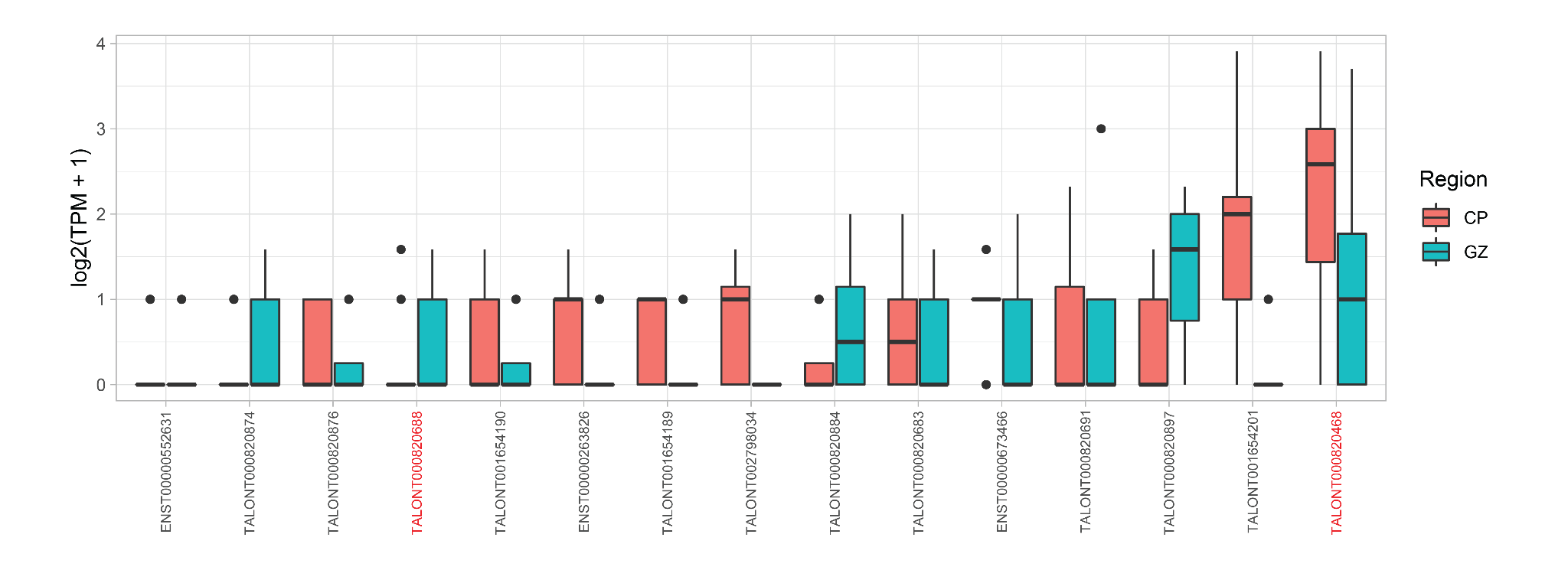

#### Fig. S14. Expression levels of detected AKT3 isoforms.

Log-transformed TPM value of detected AKT3 isoforms, stratified by cortical region (GZ/CP). Novel isoforms highlighted in Figure 6E are shown in red.

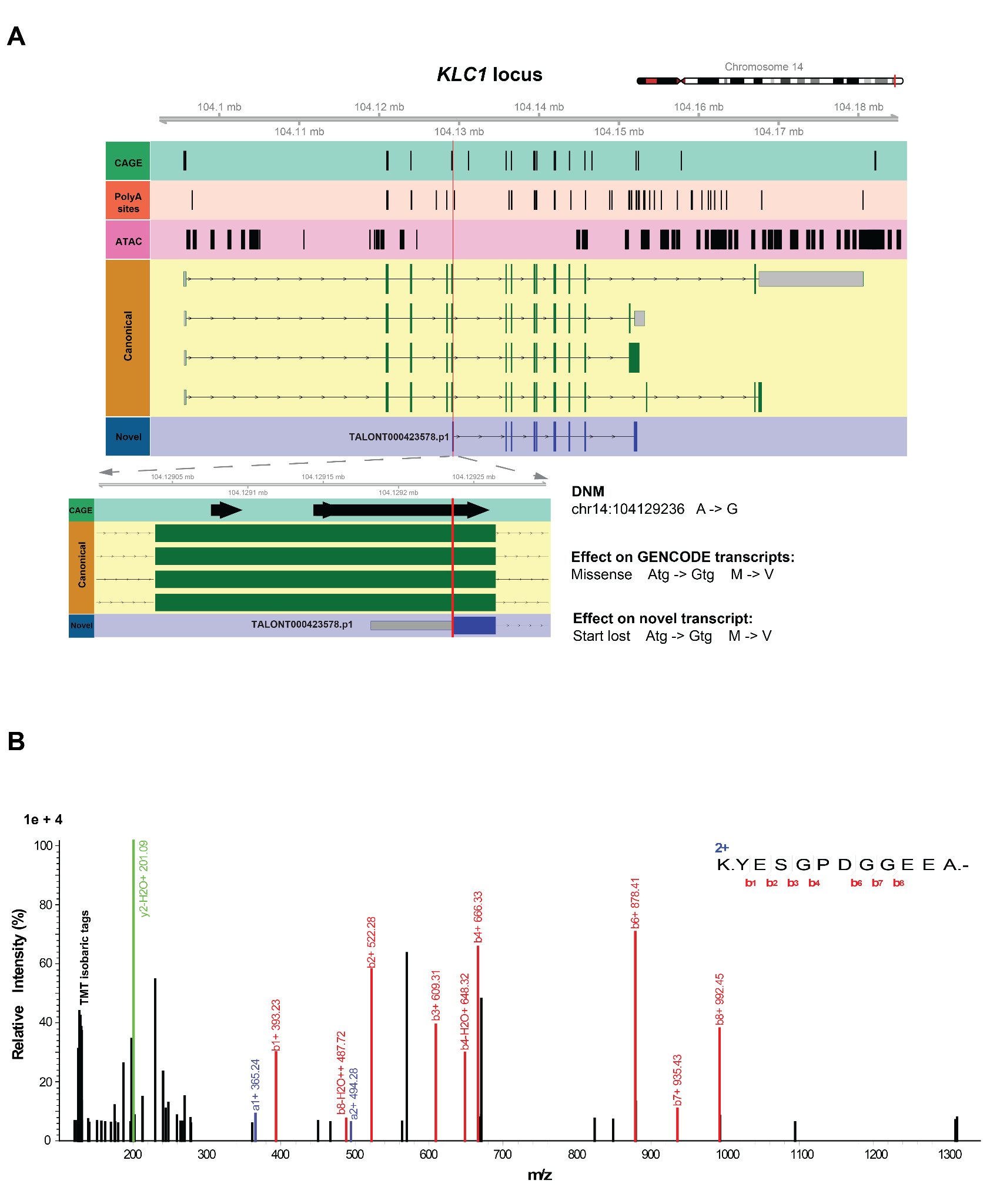

#### Fig. S15. Highly confident novel ORF explains the effect of DNM.

(**A**) The *KLC1* gene locus with representative canonical isoforms and a novel isoform identified in this study. Red vertical line indicates the position of a case DNM that affects this locus. The affected region is highlighted in the lower panel. This DNM leads to a missense mutation in the canonical protein isoforms, while causes a start lost in the novel protein isoform. (**B**) Representative mass spectrum of peptide YESGPDGGEEA at the carboxy terminus , which confirms the translation of the identified KLC1 novel protein-coding isoform. Matched b, y, and a ions were highlighted.

#### Table S1. Full-length transcriptome of the developing human brain. (A) Transcript classification using the TALON pipeline; (B) Transcript quantification using the TALON pipeline (C) Filtering criteria used to generate the TALON whitelist

#### Table S2. Transcript support from orthogonal genomic and proteomic datasets and novel exons identified (A) Novel isoforms supported by independent proteomic datasets; (B) Novel exons identified in the GZ/CP samples

#### Table S3. Summary statistics of Differential Transcript Usage during Neurogenesis. (A) Summary statistics for regional differential gene expression (DGE), transcript expression (DTE), and transcript usage (DTU) across GZ and CP . A positive log2 fold change indicates an increase in the associated gene or isoform in CP as compared to GZ. (B) Summary results and enrichment analysis of functional consequences of isoform-switch events from IsoformSwitchAnalyzeR; (C) Data support analysis of functional consequences of isoform-switch events from IsoformSwitchAnalyzeR; (D) Analysis of distal polyA site usage between GZ and CP samples

#### Table S4. Coexpression network analyses. (A) Gene- and isoform-level co-expression modules; (B) Module-level cell type enrichments; (C) Module-level rare variant enrichments; (D) Module-level RBP enrichments

#### Table S5. Single-cell analyses. (A) Cell barcodes with associated metadata and cluster labels; (B) Genomic coordinates of novel exons identified by scIso-Seq; (C) Differentially expressed isoforms across clusters; (D) Differentially expressed isoforms between pairs of clusters; (E) Differential isoform usage across clusters

#### Table S6. Genetic enrichments and variant re-interpretation. (A) Rare variant enrichment analyses among transcriptomic features (DGE/DTE/DTU genes, network modules, single cell markers). OR and P-values from logistic regression, controlling for gene-length; (B) DNMs and their most severe consequence before and after taking into account novel isoforms; (C) SpliceAI predictions for DNMs.

#### Table S7. RNA-binding protein enrichment metadata. (A) Brain-enriched RBP alternative splicing (AS) and binding (CLIP) datasets curated from literature; (B) ENCODE RBP binding (eCLIP) datasets

#### Table S8. Primers used for novel exon validation. (A) Sequences of the primers used to validate novel spliced-in exonic regions identified in this study. Primers were synthesized by Integrated DNA Technologies (IDT).

#### Data S1: This dataset contains transcriptome annotations in General Transfer Format (gtf) generated by long-read sequencing of bulk tissue samples in this study. The annotations delineate genomic coordinates, exonic/intronic boundaries, and salient attributes for each detected transcript isoform.

#### Data S2: This dataset contains transcriptome annotations in General Transfer Format (gtf) generated by long-read sequencing of single cells from human neocortex.
